# Sequential drug treatment targeting cell cycle and cell fate regulatory programs blocks non-genetic cancer evolution in acute lymphoblastic leukemia

**DOI:** 10.1101/2023.03.27.534308

**Authors:** Alena Malyukova, Mari Lahnalampi, Ton Falqués-Costa, Petri Pölönen, Mikko Sipola, Juha Mehtonen, Susanna Teppo, Johanna Viiliainen, Olli Lohi, Anna K Hagström-Andersson, Merja Heinäniemi, Olle Sangfelt

**Author notes:** Corresponding authors Olle Sangfelt, Department of Cell and Molecular Biology, Karolinska Institutet, Biomedicum, Solnavägen 9, 171 77, Stockholm, Sweden, Alena Malyukova, Department of Cell and Molecular Biology, Karolinska Institutet, Biomedicum, Solnavägen 9, 171 77, Stockholm, Sweden, Merja Heinäniemi The Institute of Biomedicine, University of Eastern Finland. Equal contribution (Alena Malyukova, Mari Lahnalampi, Merja Heinäniemi, Olle Sangfelt).

## Abstract

Targeted therapies exploiting vulnerabilities of cancer cells hold promise for improving patient outcome and reducing side-effects of chemotherapy. However, efficacy of precision therapies is limited in part because of the cellular heterogeneity of tumors. A better mechanistic understanding of how drug effect is linked to cancer cell state diversity is crucial for identifying effective combination therapies that can overcome the heterogeneity to prevent disease recurrence. Here, we characterized at the level of gene regulatory networks and at single-cell resolution the effect of G2/M cell cycle checkpoint inhibition in acute lymphoblastic leukemia (ALL) and demonstrate that WEE1 targeted therapy impinges on cell fate decision regulatory circuits. We found highest inhibition of recovery of proliferation in ALL cells with KMT2A-rearrangment (KMT2A-r), compared to cells of other leukemia subgroups. Single-cell transcriptome and chromatin accessibility profiling of (*KMT2A*::*AFF1*) RS4;11 cells treated with the WEE1 inhibitor AZD1775 revealed diversification of cell states at the fate decision points, with a fraction of cells exhibiting strong activation of p53-driven processes linked to induction of apoptosis and senescence, and disruption of a core KMT2A-RUNX1-MYC regulatory network through CDK1-mediated RUNX1 degradation. In RS4;11 cells and in patient-derived xenograft (PDX) model, we uncovered that in this cell state diversification induced by WEE1 inhibition, a subpopulation transitioned to a cell state characterized by activation of transcription factors regulating pre-B cell fate, lipid metabolism and pre-BCR signaling which supported a drug tolerance. Sequential treatment targeting the drug tolerant subpopulation with BCR-signaling inhibitors dasatinib, ibrutinib, or perturbing metabolism by fatostatin or AZD2014 after AZD1775 administration, effectively counteracted drug tolerance that drove recovery of leukemic cells. Collectively, our findings provide new insights into the tight connectivity of gene regulatory programs associated with cell cycle and cell fate regulation, and a rationale for sequential administration of WEE1 inhibitors with low toxicity inhibitors of pre-BCR signaling or metabolism.

## INTRODUCTION

The WEE1 checkpoint kinase is a key regulator of the G2/M and G1/S transitions during cell division cycle (Moiseeva, Qian et al. 2019, Elbaek, Petrosius et al. 2020) through inhibitory phosphorylation of cyclin-dependent kinases 1/2 (CDK1/2) (Parker, Sylvestre et al. 1995, Watanabe, Arai et al. 2005). WEE1 supports genome integrity by suppressing excessive origin firing during DNA replication and premature entry into mitosis (Russell and Nurse 1987, McGowan and Russell 1995, Sorensen and Syljuasen 2012). Consequently, inhibition of WEE1 removes the negative phosphorylation on CDK1/2 (in concert with CDC25) resulting in deregulation of CDKs, increased origin firing and replication fork degradation leading to fork collapse, DNA damage and unscheduled mitosis (Hirai, Iwasawa et al. 2009, Mahajan and Mahajan 2013, Moiseeva, Qian et al. 2019, Petrosius, Benada et al. 2023). Recent studies show that WEE1 inhibition by AZD1775 (Adavosertib), a potent and specific small-molecule inhibitor, alone or in combination with DNA damaging agents effectively kills cancer cells of various origins (Hirai, Iwasawa et al. 2009, Carrassa and Damia 2017, de Jong, Langendonk et al. 2019). The targeting of DNA damage response-related proteins while also challenging the cell with genotoxic agents is a widely adopted concept in cancer therapy (O’Connor 2015). However, clinical application is restrained by a high rate of toxicity in combination with chemotherapy (O’Connor 2015, Fu, Wang et al. 2018, Takebe, O’Sullivan Coyne et al. 2018, Kong and Mehanna 2021).

Acute lymphoblastic leukemia (ALL) is the most common type of childhood cancer, typically of B-cell lineage. Approximately 75% of B-ALL cases contain chromosomal rearrangements involving alteration of lineage specific transcription factors (TFs) that are critical regulators of B-cell development and serve as prognostic markers underlying clinical responses. Mixed lineage leukemia (MLL) refers to chromosomal translocations involving the gene *KMT2A* which encodes histone lysine N-methyltransferase 2, an important regulator of epigenetic maintenance of stem cell gene transcription through activating histone modification at target genomic loci (Chan and Chen 2019). Despite improved treatment protocols with higher response rates and better outcomes for children with ALL, KMT2A-r patients respond poorly to conventional chemotherapy (Liedtke and Cleary 2009, Winters and Bernt 2017). However, these leukemias typically harbor very few additional mutations that could explain drug resistance (Andersson, Ma et al. 2015). Recently, single cell genomics-approaches have provided new insight into the aberrant gene regulatory programs that cause cell state instability, and hence diversification of cell states at cell fate decision points even in isogenic cell populations in response to treatment (Pisco, Brock et al. 2013). Such non-genetic heterogeneity has emerged as a central mechanism of drug resistance in cancer, including in leukemia (Knoechel, Roderick et al. 2014, Chen, Yu et al. 2022). A network of regulatory interactions exerted by TFs poises cells at fate decision points, primed for diversification into persistent cell phenotypes. Specifically, increased drug tolerance in leukemia has been attributed to “stemness” properties, due to shift into more immature (stem-like) states-a phenotype switch governed by gene regulatory networks as opposed to resulting from genomic mutations (Ebinger, Ozdemir et al. 2016, van Galen, Hovestadt et al. 2019, Chen, Yu et al. 2022, Khabirova, Jardine et al. 2022).

Chromosomal rearrangements of the *KMT2A* gene can produce many different chimeric TFs through in-frame fusions with various partner genes (Meyer, Kowarz et al. 2009), in ALL most commonly resulting in KMT2A::AFF1 (AF4) and KMT2A::ENL fusions through t(4;11) and t(11;19) rearrangements, respectively. These fusion proteins lack the C-terminal SET domain which normally confers the KMT2A multiprotein complex histone H3K4 methyltransferase activity. KMT2A fusion proteins instead associate with the super elongation complex, resulting in differentiation arrest at the early hematopoietic progenitor state due to increased levels of activating H3K79 methylation by DOT1L at genomic loci such as *HOXA9* and *MEIS1*, which are associated with leukemic transformation (Okada, Feng et al. 2005, Winters and Bernt 2017). However, KMT2A fusions also compromise the S-phase checkpoint by abrogating stabilization of wild-type KMT2A upon DNA damage, thereby preventing trimethylation of H3K4 and causing aberrant loading of CDC45 at late replication origins (Liu, Takeda et al. 2010). Recent work further indicates that other hematopoietic master regulators of stemness and development, notably RUNX proteins, also participate in cell cycle checkpoint control and DNA repair independent of their TF activities (Friedman 2009, Wilkinson, Ballabio et al. 2013, Tay, Krishnan et al. 2018, Wray, Deltcheva et al. 2022). The impact of KMT2A fusion proteins at the convergence of two pivotal regulatory pathways, cell cycle checkpoint and cell fate control, represents a potential vulnerability; however, diversification of cell states upon targeted therapy may induce tolerance in distinct subpopulations, whose control would require combinatorial and sequential treatment.

Cancer cell responses to drug perturbations are often assayed using simple phenotypic readouts, such as proliferation or cell death. However, these end-point assays are insufficient to uncover cellular mechanisms that allow some cells to survive and that arise only in response to treatment within a subpopulation. Thus, distinguishing between selection of pre-existing drug resistant cells and drug tolerance through drug-induced mechanisms is important to predict long-term drug efficacy. Importantly, targeted therapy that impinges on core regulatory circuits affects cell fate decisions which is poised to cause cell fate diversification. The latter generates non-genetic heterogeneity that has emerged as a major driver of treatment resistance. In the present study, we characterize in detail the genome-wide response and cell state dynamics and diversification upon treatment with cell cycle checkpoint-targeted drugs in leukemic cells. We used single cell resolution multi-omics and demonstrate that perturbing the cell cycle regulation directly impacts cell fate regulatory circuits and cell fate decision, resulting in transition to a drug-tolerant cell phenotype in a particular subpopulation. Contrary to currently held views that stemness features attributed to KMT2A-leukemias critically underlie drug resistance, we found that upon targeting the cell cycle control vulnerability of KMT2A-r leukemias, the drug-tolerant cells corresponded to a more differentiated phenotype with expression of pre-BCR and BCL6. We show that WEE1 inhibition followed by drugs targeting the persisting cell state regulatory program prevents recovery in KMT2A-r and pre-BCR+ ALL cells. This synergy was observed only in the sequential, not in concurrent treatment schedule.

## RESULTS

### WEE1 inhibitor AZD1775 results in prolonged growth-inhibition selectively in KMT2A-r B-ALLs

*WEE1* expression was previously reported to be higher in primary ALL blasts compared to normal mononuclear cells (Ghelli Luserna Di Rorà, Beeharry et al. 2018). To determine whether specific leukemia subtypes may rely more on *WEE1* expression, we used our data resource Hemap (http://hemap.uta.fi/) which contains curated genome-wide transcriptomic data from more than 30 hematologic malignancies. Interestingly, *WEE1* expression was significantly higher (adjusted p-value <0.01, Wilcoxon test) in the KMT2A-r ALL subtype (Figure 1A, see also Figure S1A comparing KMT2A::AFF1, KMT2A::MLLT1 and KMT2A::MLLT3). Higher *WEE1* expression in KMT2A-r compared to other ALL subtypes raised the possibility that KMT2A-r cells may be more vulnerable to targeted inhibition of the WEE1 kinase. However, since WEE1 activity is regulated also at the protein level via phosphorylation, we performed further functional evaluation comparing different leukemic cell lines. First, to assess sensitivity of leukemic cells that represent different ALL subtypes and lymphoid lineage differentiation states to WEE1 kinase inhibition, we treated a panel of 15 ALL cell lines with increasing concentrations of AZD1775 (Figure 1B). Acute drug treatment induced apoptosis as measured by activation of caspase-3/7 and reduced cell viability in a dose-dependent manner in all cell lines (Figure 1B, Figure S1B).

**Figure 1.**
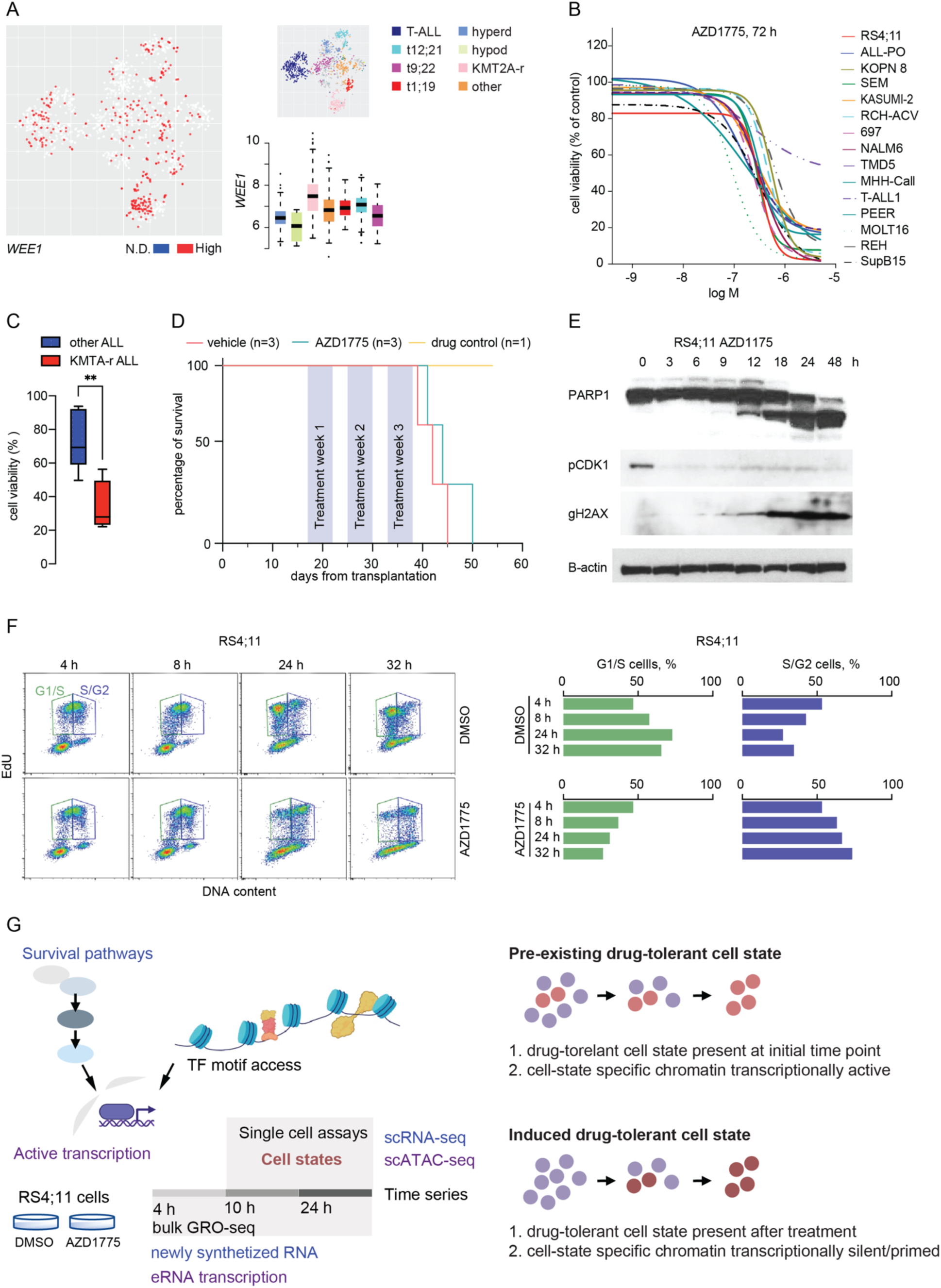
AZD1775 response profiling reveals dependency on WEE1 in KTM2A-r ALLs *in vitro* and emergence of drug tolerance in a monotherapy setting *in vivo*. **A.** *WEE1* gene expression in primary ALL samples in HEMAP. tSNE-map: low (in white) vs high (in red) mRNA level is visualized for each patient transcriptome profile. Boxplot: normalized gene expression level comparing across subtypes. Subtype annotation is indicated by different colors (right panel). **B.** AZD1775 dose response profiles (relative to DMSO control) were assessed by Alamar Blue assay in T-ALL1, Molt16, Peer, RCH-ACV, 697, Kasumi-2, MHH-Call3, SupB15, TMD6, Nalm-6, REH, RS4;11, SEM and KOPN8 ALL cells. Percentage of viable cells after 72 h treatment with increasing concentrations of AZD1775 is shown. **C.** Recovery of proliferation following removal of AZD1775 (relative to DMSO) analysed by Alamar Blue assay. Cell lines were treated for three days with AZD1775 (corresponding IC50 for each cell line, (see Fig S1B) and allowed to recover for an additional 14 days without the drug. Box plot comparing recovery in KMT2A-r and non-KMT2A-r ALL cell lines (corresponding subtypes TCF3-PBX1, ETV6-RUNX1, TCF3-PDGFRB) is shown. ** denotes p=0.0086 determined using two-tailed t-test. Data are represented as mean ± SD. **D.** Kaplan-Meier survival curves of MLL-7 patient-derived KMT2A-r cells treated in a xenograft model with AZD1775. **E.** Cellular response to AZD1775 in KMT2A-r RS4;11 cells was assessed at the indicated time points by Western blot analysis. Whole cell lysates were immunoblotted with the specified antibodies; Cleaved PARP1 (marker of cell death), phosphorylation of CDK1-Y15 (marker of WEE1 inhibition) and phosphorylation of γH2AX-S139 (marker for DNA damage). β-actin was used as a loading control. **F.** Flow cytometry density plots showing distribution of EdU (replicating cells, Y-axes) and DNA content (Propidium Iodide, PI, X-axes) to determine the proportion of cells in G1/S (green) and S/G2M (purple). RS4;11 cells were pulse-labelled for 30 minutes with EdU, treated with DMSO or AZD1775 and chased for the times indicated. **G.** Overview of the experimental setup for genomics characterization of early drug response dynamics using GRO-, scRNA-, and scATAC-seq to distinguish pre-existing and induced drug tolerance. Time series (4, 10 and 24 h treatment) in GRO-seq coupled with cellular resolution single cell genomics assays at 10 and 24 h to detect drug-tolerant cell states and decipher their corresponding survival pathway and regulatory network activity.

Next, we assessed whether the response to AZD1775 persisted, by measuring recovery of proliferation post-treatment in leukemia cell lines, comparing the different chromosomal translocations, including *ETV6-PDGFRB* (Nalm-6), *TCF3-PBX1* (697, RCH-ACV and Kasumi-2), *ETV6/RUNX1* (REH) and KMT2Ar-ALLs (RS4;11, SEM, ALL-PO and KOPN8). Cells were treated with AZD1775 at sublethal dose (GI50 concentration) for three days, followed by drug washout and regrowth in drug-free media. Although acute response to AZD1775 (GI50) did not associate with any specific ALL subtype (Figure S1B), WEE1 inhibition markedly attenuated re-proliferation following drug washout in KMT2A-r cells as opposed to non-KMT2A-r cells (Figure 1C, Figure S1D). Thus, the KMT2A-r subtype with the highest expression of *WEE1* exhibits significantly reduced recovery, in line with the patient genomics cohort (Figure 1A) result. To provide independent cell line evidence based on Crispr-Cas9 targeting across haematopoietic malignancies, we further analyzed *WEE1* gene dependency scores using the DepMap portal (https://depmap.org/portal/) and found a greater dependency on WEE1 in leukemia cell lines with higher *WEE1* expression (Spearman r =-0.336, Figure S1C). However, characterization of the response to AZD1775 monotherapy cycles *in vivo*, using a primary KMT2A-r patient-derived xenograft (PDX) model MLL-7 (*KMT2A::AFF1*) in (Richmond, Carol et al. 2015) NOD.Cg-*PrkdcscidIl2rgtm1Wjl*/SzJ (NSG) mice (N=3 per group) transplanted with leukemic cells did not indicate a strong survival benefit (Figure 1D, AZD1775 median survival; 44 days, DMSO median survival; 42 days, see also Figure S1E-I), warranting a therapy strategy informed by cellular properties that contribute to drug tolerance.

To explore the molecular determinants of the AZD1775 response, we selected KMT2A-r RS4;11 cells and non-MLL-r Nalm-6 cells for more in-depth analysis. First, we confirmed that AZD1775 reduced phosphorylation of CDK1 on threonine 15, increased PARP cleavage and triggered DNA damage rapidly, as measured by phosphorylation of γH2AX (Figure 1E, RS4;11 cells), according to the established mode of action of AZD1775. To confirm that RS4;11 cells treated with AZD1775 enter G2/M prematurely, cells were pulse-labelled with EdU and treated with DMSO or AZD1775 for different times, up to 32 hours. As shown in Figure 1F, while DMSO-treated cells divided and re-entered G1-phase, a large proportion of AZD1775 treated RS4;11 cells stalled at G2/M and failed to proceed through the cell cycle.

Since the KMT2A-r cell lines tested retain the KMT2A-fusion and represent a genetically homogenous clonal origin, we asked whether drug tolerance in the recovering cells is derived from (1) a pre-existing drug-tolerant cell subpopulation that are selected for and expand or is (2) acquired during treatment through drug-induced cell state transition (Cohen, Geva-Zatorsky et al. 2008, Sharma, Lee et al. 2010, Pisco, Brock et al. 2013). The latter would involve a shift of gene expression program, and thus would be driven by TF activity changes whereby possibly only a fraction of cells might be capable to accommodate a new phenotypic state that supports drug tolerance. To distinguish between these two schemes, we used the RS4;11 cells as a cellular model system (Figure 1G). We first performed global run-on sequencing (GRO-seq) to assess early effects on newly synthesized RNA at gene and enhancer regions at 4, 10 and 24 hours. Changes in the pattern of ongoing transcription following drug treatment would support the second possibility if it can be observed in a substantial fraction of cells and already at an early time point before the first cell division. The 10 and 24 h time points were then further resolved using single cell RNA (scRNA)-and ATAC (scATAC)-sequencing to the determine the fraction of cells that exhibit the putative tolerance conferring transcriptional response at the level of transcription and chromatin access. We then used the combination of these assays to inform about the presence of drug-tolerant cell states, association between early transcriptional changes, chromatin access and regulatory network activities (Figure 1G).

### WEE1 kinase inhibition triggers a rapid transcriptional response and emergence of multiple cell states

The time course GRO-seq profiles identified 1239 significant gene transcription changes comparing across time points (F-test adjusted p-value < 0.001). The temporal response to AZD1775 in bulk (Figure 2A) could distinguish: 1) genes primarily upregulated at 4 hours, exemplified by the GRO-seq signal at the *PLK1* gene locus (Figure 2B), 2) genes peaking in upregulation at 10 hours, 3) genes most strongly upregulated at 24 hours, and 4) genes downregulated at all time points (see also Table S1). Pathway enrichment analysis revealed genes associated with regulation of the cell cycle, DNA replication, PLK1/AURORA signaling, FOXM1 TF network and M-phase pathways in the early (4 h) upregulated cluster (Figure 2C). Genes functionally related to cell cycle checkpoints, DNA-strand elongation, activation of pre-replication complexes and S-phase were repressed in the bulk genomics data (Figure 2A bottom, “down”).

**Figure 2.**
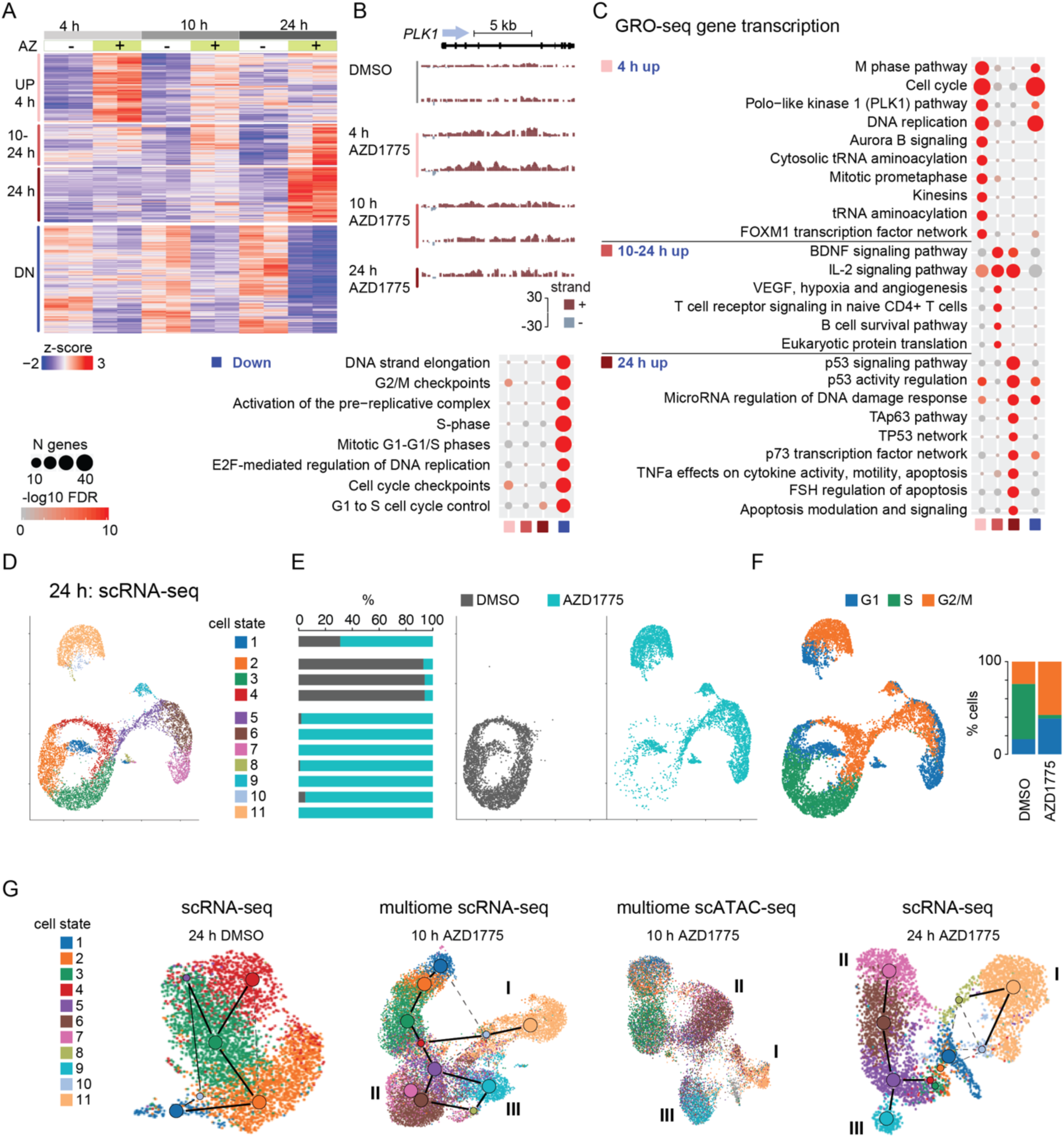
WEE1 kinase inhibition triggers a rapid transcriptional response and emergence of multiple cell states. **A.** Heatmap illustrating the magnitude and direction of changes in the GRO-seq signal (z-score; tones of red indicate high level, tones of blue low level) for annotated gene transcripts. Gene clusters that correspond to regulation patterns 4 h up, 10-24 h up, 24 h up, and down are shown. **B.** GRO-seq signal is illustrated at *PLK1* gene region from replicate samples collected at 4, 10 and 24 h (top left). Signal from +/-stand is plotted above (in red) or below (in grey), respectively. **C.** Enrichment of biological processes for GRO-seq gene clusters (right: upregulated with distinct time profile, left: downregulated) shown as dot plots. Size of the dot corresponds to the number of genes from the gene set and-log10 FDR value is shown in color. **D-F**. Low dimensional projection and scRNA-seq transcriptome-based clustering for the RS4;11 cells from DMSO and AZD1775 treated cells are shown on the UMAP visualization. Colors correspond to cell state assignment (in D), treatment (in E), and assigned cell cycle status (F). Quantification of cell proportions across categories is shown in E and F. **G**. Low dimensional projections based on transcriptome and chromatin access profiles are shown in UMAPs generated separately for DMSO and AZD1175 treated cells. Graph connectivity (PAGA) is visualized on scRNA-seq maps where connecting lines indicate putative cell state transition paths. Multiome data from 10 h includes scATAC-seq from the same cells. The colors correspond to assigned cell state based on 24 h scRNA-seq data, refer to cell states shown in D. Three treatment-specific cell fates that are reproducibly found in both time points and data modalities are indicated by roman numerals (cell fate I, II and III).

Since the WEE1 inhibition interferes with a central decision point of cell fate control during cell cycle, we anticipated that this perturbation would result in cells occupying distinct states. To expose the diversification and nature of responses of individual cells we therefore performed single-cell transcriptome and chromatin access measurements. scRNA-sequencing revealed that 24 h after treatment with AZD1775, the surviving cells analyzed alongside DMSO treated cells formed 11 discernable cell clusters (Figure 2D). We refer to these clusters as cell states hereafter. As determined by expression of cell cycle phase specific markers, the clusters 2, 3 and 4, correspond to different cell cycle phases present in DMSO-treated cells. The other clusters, 5-11, were almost completely represented by AZD1775 treated cells (Figure 2E). As noted in the bulk GRO-seq analysis, treated and surviving cells had reduced number of cells in the transcriptional state associated with S phase and an enrichment of cells in G1 or G2/M phase (Figure 2F), indicating an arrest at the G1/S checkpoint and a possible failure to progress through mitosis, consistent with the flow cytometry data of Figure 1F.

Since the bulk GRO analysis indicated ongoing transition and possible adaptation of cells arising already at 10 h, we performed single-cell RNA-and ATAC-seq cells based on the multiome protocol to assay simultaneously from the same cell how open chromatin sites impact the transcriptional response and cell state diversity, and also compared the cell states to the scATAC-seq profile acquired at 24 h from parallel cell cultures.

First, to investigate the correspondence between cell states identified from the single cell transcriptome and clustering of cells defined by their chromatin access, we examined the transcriptome and chromatin dynamics by separating the cells by treatment from 10 and 24 h (Figure 2G, Figure S2). Matching cell clustering could be distinguished from the separate clustering of multiome scRNA-and scATAC-seq data and between the 10 and 24 h time points (transcriptome-based cell state labels from 24 h scRNA-seq are shown, see also Figure S2B-G).

Secondly, we contrasted the alternative scenarios of differential survival of pre-existing cell states and emergence of treatment-induced cell states. The predominant cell state dynamics (Figure 2G left) in DMSO treated cells matched to active cell cycle transitions, as indicated by lines on the UMAP based on RNA velocity and PAGA graph analysis (see Methods). By contrast, AZD1775 treated cells at 10 and 24 h (Figure 2G, middle and right, respectively) followed a trajectory along three distinct branches. The first branch is dominated by cell state 11 (cell fate I), a second branch corresponds to a succession of cell states from 5 to 7 (cell fate II), and a third small population of cells branch to cell state 9 (cell fate III) (Figure 2G), each trajectory being unique to AZD1775 treated cells.

Altogether, despite the highly heterogeneous response, we found high concordance between the cell subpopulations analyzed using both RNA-and ATAC-seq analysis, and between bulk and single cell pathway analyses. This initial analysis supports the scenario that AZD1775 treatment could impact the gene regulatory network, leading to the emergence of new cell states with unique active transcription profiles. Next, we proceeded to elucidate whether this grouping based on the transcriptome and chromatin dynamics indeed reflects functional cell fates based on additional analyses.

### Forced mitotic entry in response to AZD1775 treatment arrests cells from S-phase in a condensed mitotic-like chromatin state

Cell state 11 distinguished a distinct subpopulation (cell fate I) that was evident already after 10 h and persisted at 24 h of AZD1775 treatment (Figure 2G middle and right, respectively). Based on scATAC-seq TF motif analysis, these cells had elevated S-phase (E2Fs, YY1, Table S3) and mitotic checkpoint (NRF1, Ronin, Sp1) TF motif access (Figure 3A and Figure S3A,B). Simultaneously, these cells showed evidence of mitotic entry downstream the forced activation of CDK1 based on the scATAC-seq fragment length distribution, with cells corresponding to cell fate I exhibiting a condensed chromatin state and elevated nucleosome signal metric (Figure 3B, see also Figure S3C,D). Moreover, pathway enrichment from scRNA-seq cell cycle phase marker genes was consistent with a mitotic-like state (Figure S2H and Table S2). Premature entry into mitosis can extend mitosis and enhance cell death (Visconti, Della Monica et al. 2015). In agreement with delayed mitotic exit, the fraction of phosphorylated serine 10 histone 3 (pS10-H3) with 4N DNA content increased with treatment time, reflecting a mitotic-like arrest (Figure 3C).

**Figure 3.**
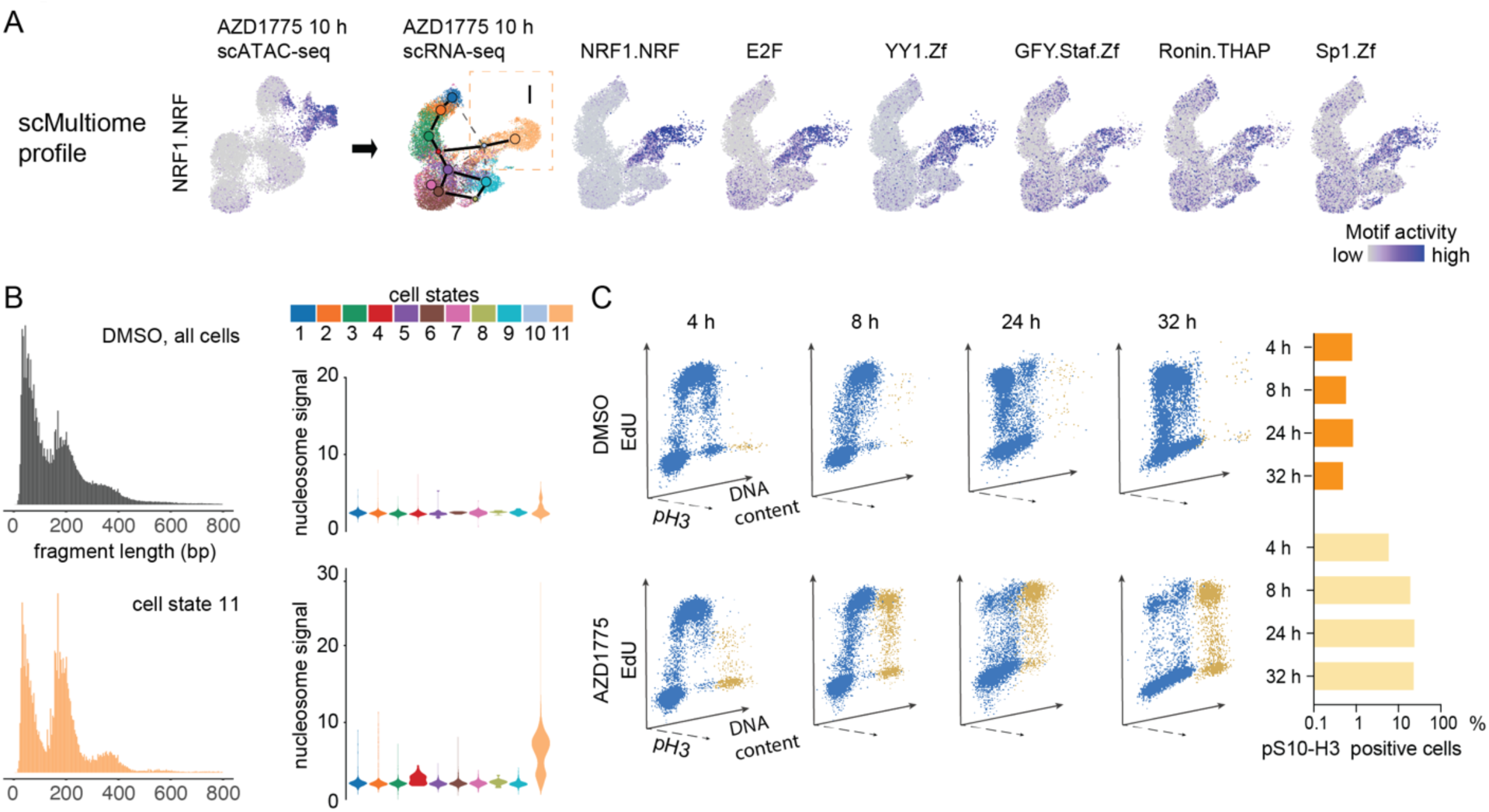
Forced mitotic entry in response to AZD1775 treatment arrests cells from S-phase in a condensed mitotic-like chromatin state. **A.** TF motif activity (ChromVar) is shown for NRF1, E2F, YY1, GFY, Ronin and Sp1 with highest motif access in cell fate I. To match with scRNA-seq cells states, TF motif activities are visualized on the 10 h scRNA-seq UMAP from the 10 h multiome profile. NRF1 motif with highest cluster-specific activity (summarized in Table S3) is shown on multiome scATAC-seq UMAP (10 h). Darker tones correspond to high motif access. **B.** Typical scATAC DNA fragment length distribution in RS4;11 cells (24 h DMSO-treated cells, top left) compared to cells assigned to cell state 11 (bottom left). The nucleosome signal metric calculated from the ratio of fragments between 147 bp and 294 bp (mononucleosome) to fragments < 147 bp (nucleosome-free) across cell states is shown as violin plots (right panel, top: DMSO, bottom AZD1775-treated cells). **C.** Three-dimensional (3D) flow cytometry dot plots showing the distribution of EdU (replicating cells, Y-axes) versus DNA content (X-axes). RS4;11 cells were pulse-labelled for 30 minutes with EdU, treated with DMSO or AZD1775 and chased for the times indicated. Mitotic status was analysed by labelling of phosphorylated S10-H3 (pH3) at the indicated time points (Z-axes). Graphs (right panel) represent the proportion of pS10-H3 positive cells in DMSO or AZD1775-treated cells.

We also performed scRNA-seq in non-KMT2A-r Nalm-6 cells for comparison. In sharp contrast to RS4;11, the majority of AZD1775 treated Nalm-6 cells retained a population in the active cell cycle (matched to cell states 2-4), in line with their ability to rapidly recover and proliferate upon drug withdrawal (Figure S3E). Accordingly, phosphorylation of S10-H3 was only transiently increased in Nalm-6 cells, while it was continuously high in RS4;11 cells treated with AZD1775 (Figure S3F), supporting a reduced capacity of KMT2A-r cells to cope with AZD1775-driven CDK-activation and replication stress compared to non-KMTA2-r cells.

**AZD1775 triggers unscheduled replication characterized by genome-wide p53 response** Comparison of the other cell clusters detected from scATAC-seq (referred to as “chromatin states”) revealed that AZD1775-induced genome-wide changes in chromatin access with overall highly distinct TF motif access patterns (Figure 4A) that could underlie the emergence of distinct cell fates upon replication stress.

**Figure 4.**
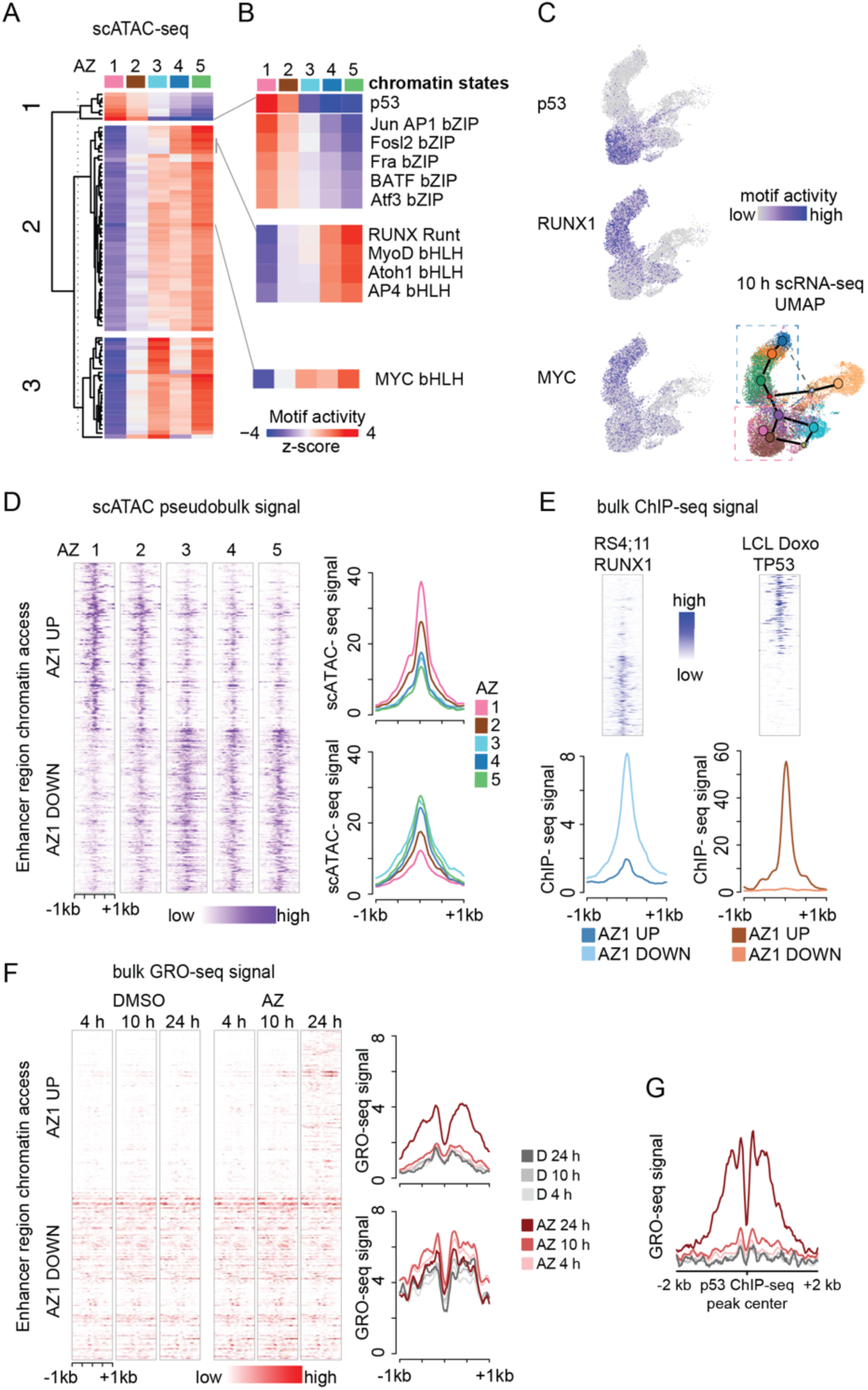
p53-driven gene regulatory response initiates from transcriptional silent chromatin and anti-correlates with RUNX1 binding. **A.** TF motif activity (scATAC, 24 h) across chromatin states 1-5 in AZD1775 treated cells is shown as a heatmap, where TF motifs (in rows) are clustered into three main patterns. **B.** Pattern 1 TF motifs with highest motif access in cell fate II (top panel). Selected pattern 2 TFs are shown for comparison (bottom panel). **C.** p53, RUNX1 and MYC TF motif activity (10 h) visualized on UMAPs, as in Fig. 3A. **D.** Pseudobulk scATAC-seq signal at-/+ 1kb at 200 up-and downregulated enhancer regions most specific to AZD1775 chromatin state 1 (AZ1) vs other cell states. In the heatmap (left) each row corresponds to one enhancer and brighter color tones to higher access level. The magnitude and direction of changes in the cell state-specific chromatin accessibility signal across chromatin states 1-5 are summarized as average signal histograms (right). **E.** ChIP-seq signal profile of RUNX1 in RS4;11 cells (DMSO) and p53 from doxorubicin (Doxo) stimulated lymphoblastoid cells (LCL). TF occupancy signal heatmap and histogram at the same enhancer regions as shown in 4D. **F.** GRO-seq signal heatmap (left) and histogram (right) at the same enhancer regions as shown in 4D. Line color corresponds to treatment time and condition. **G.** GRO-seq signal histogram at ChIP-seq-based high-occupancy p53 binding sites from LCL shown as in 4F.

Elevated access at binding motifs for p53 (and bZIP-factors JUN/AP, FOS, Atf) was characteristic of AZD1775-specific chromatin states 1 and 2 (Figure 4B). As can be seen from the TF motif access score visualized on the scRNA-seq map (Figure 4C), these cells corresponded to the branch leading to cell states 6-7 on the scRNA-seq map (cell fate II) (Figure 4C).

Based on the scATAC-seq profiles of the distinct cell states, the elevated p53 TF motif access (Figure 4A, in pattern 1) anti-correlated with two other TF motif access patterns (Figure 4A, patterns 2 and 3). Closer inspection of pattern 2 revealed mutual exclusivity with the lymphoid progenitor TF RUNX1 (clustered together with bHLH motifs recognized e.g. by AP4) (Figure 4B, Figure S5A). RUNX1 is a key TF regulating target genes downstream of KMT2A-AFF1 in a feedforward loop (Harman, Thorne et al. 2021) and was expressed at higher levels in KMT2A-r patients compared to other ALL subtypes (Figure S5A). To further explore the relationship between cell state-specific chromatin accessibility and transcriptional regulation, we visualized the scATAC-seq signal profile at enhancer regions with highest or lowest chromatin access in chromatin state 1 (Figure 4D, AZD1775-treatment). Next, we generated RUNX1 ChIP-seq profile from RS4;11 cells (basal state) and retrieved p53 ChIP-seq signals from doxorubicin stimulated lymphoblastoid cells (Figure 4E). The average TF occupancy signals at these cell fate II-specific high chromatin access regions showed high p53 and low RUNX1 signal, while an opposite profile characterized the low access regions, providing further confirmation of their mutually exclusive TF activities. Notably, cells with high RUNX1 TF motif activity in AZD1775-treated cells (chromatin states 4 and 5) corresponded to transcriptome cell states 1-4 (present also at basal state), that in relative cell proportion decreased from 60% at 10 h to 11% at 24 h.

To further explore the relationship between cell state-specific chromatin accessibility and transcriptional regulation, we quantified transcription from enhancers (enhancer RNA, eRNA) at these same regions (AZD1775, chromatin state 1). This regulatory region activity profile showed that enhancers in cell fate II-specific accessible chromatin were transcriptionally silent at 4 and 10 h (Figure 4F). This is in agreement with gene-level GRO-seq results that indicated delayed upregulation of the p53-signaling network and pathways regulating apoptosis (cluster active at 24 h, Figure 2A). Enrichment of p53 and bZIP binding motifs in enhancer regions active at 24 h was independently validated in the bulk GRO-seq data (see Methods, Figure 4G Figure S4D, Table S4). Hence, inducing a p53 response from initially silent regulatory regions could allow cells time to adapt and initiate pro-survival responses.

### CDK-mediated degradation results in loss of RUNX1 in response to AZD1775

To further characterize the decline in RUNX1 activity and other pattern 2 TFs, including the KMT2A::AFF1 TF target MYC that promotes leukemia survival (Harman, Thorne et al. 2021) we performed immunoblot analysis (in bulk) and found that AZD1775 treatment decreased RUNX1 protein expression levels in KMT2A-r RS4;11 and SEM cells, and in several non-KMT2A-r ALL cell lines (Figure 5A, Figure S5B). The hematopoietic stem cell TF GATA2 and MYC similarly had declining protein expression following AZD1775 treatment (Figure S5D-E). In comparison, ATF4 that acts as transcriptional activator of the integrated stress response was transiently induced at 4 h (Figure S5C).

**Figure 5.**
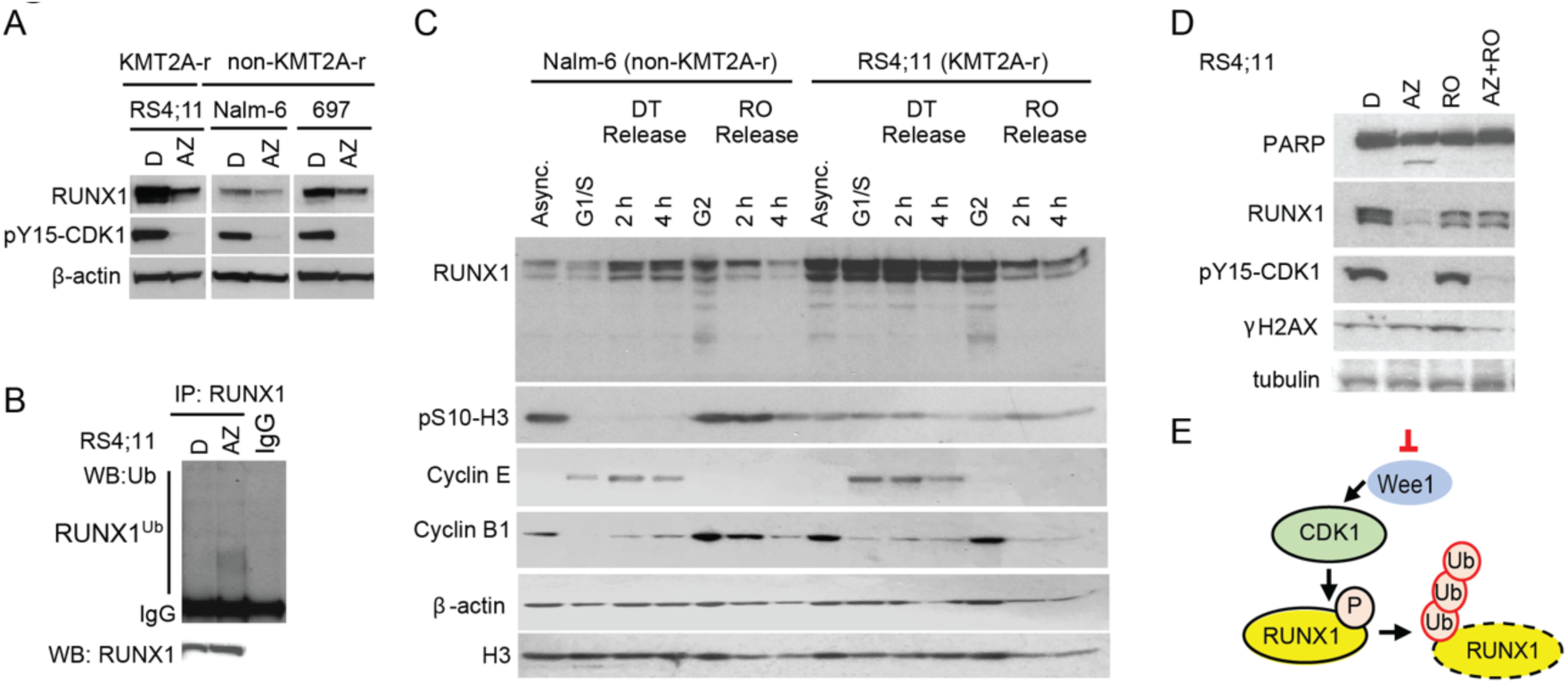
WEE1 inhibition promotes CDK-driven RUNX1 protein degradation. **A.** RUNX1 protein levels in RS4;11, Nalm-6 and 697 treated with AZD1775 or DMSO for 24 h analysed by immunoblotting. Phosphorylation CDK1 was detected with pY15-CDK1 antibodies. β-actin is a loading control. **B.** Poly-ubiquitination of RUNX1 in RS4;11 cells treated with AZD1775 for 24 hrs (and with proteasomal inhibitor MG-132 for the last 3 hr). RUNX1 was immunoprecipitated (IP) under denaturing condition to disrupt non-covalently linked ubiquitin. RUNX1-ubiquitination was detected by immunoblotting for ubiquitin antibodies. 10 % of whole cell lysate was used as input control for IP. **C.** RS4;11 and Nalm-6 cells were synchronized at different cell cycle stages (G1/S or G2) by double thymidine block (DT) or by treatment with RO3306 (RO) (CDK1 inhibitor), and release for the indicated time points. RUNX1 protein levels were analysed in by immunoblotting of whole cell lysates using RUNX1 antibodies. Asynchronous (Async) cells were treated with DMSO. Expression of cyclin E, cyclin B1 and phosphorylation of pS10-H3 was analysed as indicated. β-actin and H3 are loading controls. **D.** RS4;11 cells treated with AZD1775 (AZ), RO3306 (RO) or their combination (AZ + RO) for 24h. Expression of RUNX1, cleaved PARP1, phosphorylation of Y15-CDK1 and phosphorylation of S139-γH2AX was analyzed by immunoblotting of whole cell lysates with the antibodies as indicated. Tubulin is a loading control. **E.** Schematic illustration of mechanism linking CDK1 activity and RUNX1 protein degradation following WEE1 inhibition by AZD1775.

RUNX1 was previously reported to be degraded in a CDK-dependent manner by the APC ubiquitin ligase complex during mitosis (Biggs, Zhang et al. 2005, Wang, Zhang et al. 2007). Based on the scATAC-seq TF motif analysis in DMSO-treated cells, the RUNX1 and bHLH motifs have the highest access at G1 and lowest access in G2 (Figure S5D, see also Figure S2C). We hypothesized that without the inhibitory phosphorylation by WEE1 (AZD1775 treatment), hyperactivation of CDK1/2 may promote ubiquitination and proteasomal degradation of RUNX1 during G2/M. Supporting this, AZD1775 treatment did not alter *RUNX1* mRNA expression levels in RS4;11 cells or in other B-ALL cell lines (Figure S5E), but increased RUNX1 protein ubiquitination upon proteasomal inhibition (Figure 5B). We next verified that RUNX1 levels decreased following release from a G2/M-phase block as compared to cells released from a G1/S-phase block (Figure 5C). Concurrent treatment with the CDK1 inhibitor RO3306 and AZD1775 partially rescued downregulation of RUNX1 protein, and reduced PARP cleavage and γH2AX (Figure 5D), further supporting CDK1-driven ubiquitination and proteasomal degradation of RUNX1 in response to WEE1 inhibition (illustrated in Figure 5E).

Together, these results suggest that WEE1 inhibition may interrupt a core KMT2A-r transcriptional regulatory program involving RUNX1-MYC-GATA TFs through reduced protein expression.

### Stress-and pre-B-state regulatory programs distinguish a drug-tolerant sub-population with pre-BCR and BCL6 gene loci activation

Functionally, RUNX1 is part of a core regulatory circuit that acts as a switch, governing cell fate commitment (Harman, Thorne et al. 2021). Consequently, perturbing RUNX1 expression and TF activity dynamics by WEE1 inhibition may induce a transition to an alternative cell state. In line with this, the cell fate III subpopulation (cell state 9, Figure 2G) had decreased RUNX1 motif activity (Figure 4A, AZD chromatin state 3), but significantly higher chromatin access of pre-B cell state TF motifs, including PAX5, EBF, GR and MEF2C, and high motif activity of TFs regulating cellular metabolism (SREBF, THR, LXR) (Figure 6A, see also Figure S6A-B). As a distinct feature, cell state 9 (cell fate III)-specific open chromatin regions had highest NfKB and heat shock factor (HSF) motif access (Figure 6A). Supporting activation of a functional TF circuitry, cell fate III cells had high expression of the respective TF target genes of SREBF (*LDLR*, *HMGCS1*), HSF (*HSPA1B*), GR (*TSC22D3*), NFkB (target gene set activity scores shown) and pre-BCR signaling (*BCL6, SOCS1)* (Figure S6C-E, Table S2). The combination of stress-and pre-B fate specific TF activation pointed towards cell state transition to a more tolerant cell state in response to AZD1775 treatment. In a similar fashion as examining cell fate II-specific chromatin (Figure 4D-F), we quantified enhancer regions that represent highest vs lowest sc-ATAC-seq signal in cell fate III (AZ chromatin cluster 3) and compared chromatin access (Figure 6B, scATAC-seq signal) and enhancer activity (Figure 6C, GRO-seq signal). Interestingly, the regions with high access in cell fate III were transcribed, although at low level, also in basal state (grey lines, Figure 6C), indicative of a primed state. Upon AZD1775 treatment, these enhancers were rapidly further activated (4 and 10 h profiles in red, Figure 6C). Comparison to NfKB ChIP-seq signal in lymphoblastoid cells provided further confirmation that the activated chromatin regions in cell fate III-specific open chromatin harbor NFkB binding sites (Figure 6D, upper panel). In comparison, the ChIP-seq signal at enhancers with lowest chromatin access revealed that these sites were p53-bound in lymphoblastoid cells treated with doxorubicin (Figure 6D, lower panel), which could indicate active suppression of p53 chromatin recruitment in cell fate III. Moreover, chromatin access was increased at gene loci encoding *MEF2D, SREBF1, BCL6, SOCS1* and pre-BCR signaling pathway components that are central to cellular survival pathways in B-lymphoid cells (Figure 6E), resembling a previously recognized leukemia-associated TF circuitry active in MEF2-fusion subtype of ALL (Tsuzuki, Yasuda et al. 2020). Consistent with induced drug-tolerant state arising through transcriptional regulation, survival pathway activation was significantly enriched at the 10-24 h activated gene cluster identified from bulk GRO-seq profile (Figure 2C).

**Figure 6.**
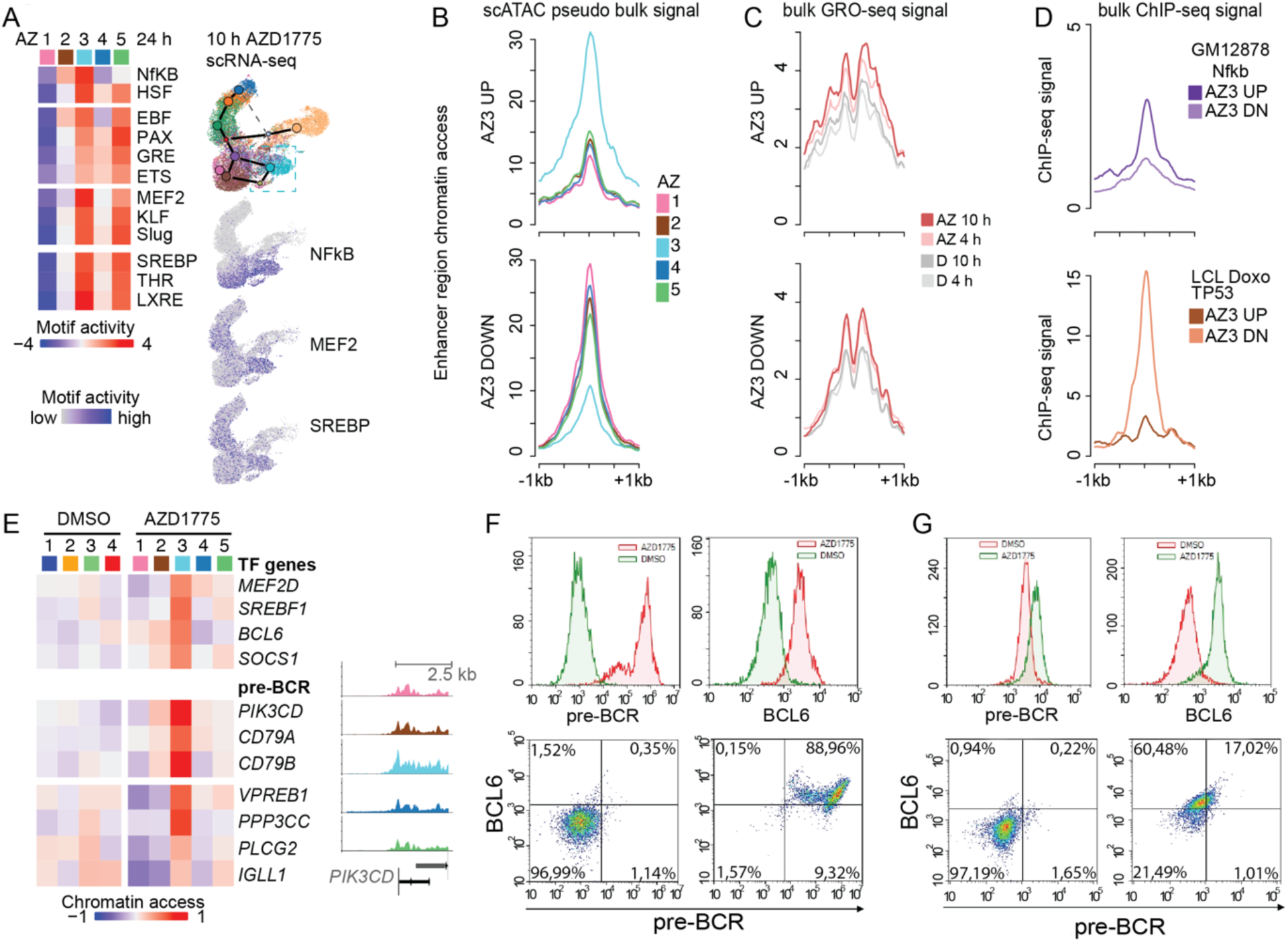
Stress-and pre-B-state regulatory programs distinguish a drug-resistant sub-population with pre-BCR and BCL6 gene loci activation. **A.** TF motif access across AZD1775 chromatin states (1-5) is shown for TFs with high activity in cell fate III (Pattern 3, see Fig. 4A) (left). Nf-KB, MEF2 and SREBP motif activity are visualized on the 10 h scRNA-seq UMAP (right). **B.** scATAC-seq signal at-/+ 1kb at 200 up-and downregulated enhancer regions most specific to AZD1775 chromatin state 3 (AZ3) vs other cell states is shown as average signal histogram, as in Fig. 4D. **C.** GRO-seq signal histogram at the same enhancer regions as shown in 6B. **D.** ChIP-seq signal profile of NfKB in lymphoblastoid cells (LCL) and p53 from doxorubicin (Doxo) stimulated lymphoblastoid cells. TF occupancy signal histogram is shown at the same enhancer regions as shown in 6B. **E.** Chromatin access at TF and pre-BCR signaling related gene loci (gene body flanked by 2.5 kb up-and downstream) compared across AZD1775 chromatin states is shown as a heatmap (left). Aggregated scATAC-signal across cells assigned to each cluster is exemplified at *PIK3CD* gene region (right). The track colors correspond to 24 h AZD1775 chromatin state annotation. **F.** Representative flow cytometry histograms (upper panels) and dot plots (lower panels) of pre-BCR receptor and BCL6 in RS4;11 cells treated with DMSO (red) and AZD1775 (green) for 72 hours (F) and following 10 days drug washout (G).

To investigate whether the emergence of cell fate III constitutes a potential escape mechanism and drug-tolerant cell state linked to recovery following AZD1775 treatment, we analysed pre-BCR and BCL6 protein expression in cells treated with AZD1775 for 72 h, and after 10 days of recovery. Notably, the drug tolerant cells that persisted at 72 h matched the pre-BCR+/BCL6+ population (cell fate III). At 72 h, pre-BCR levels were strongly upregulated in RS4;11 cells (Figure 6F top panel). This activation occurred in a reversible manner, as pre-BCR expression was restored to basal levels after 10 days recovery (Figure 6G). By contrast, BCL6 protein levels increased at 72 hours and the expression persisted for 10 days (Figure 6F,G). Based on the data obtained, we hypothesized that leukemic cells that can readily adopt the BCL6+pre-BCR+ cell state may have higher tolerance to AZD1775. The difference in the temporal dynamics of transcriptional activation of cell state II vs cell state III enhancers observed in RS4;11 cells, together with the lower chromatin access at p53 binding sites, could increase drug tolerance in cell fate III. The Nalm-6 cells used as comparison represent a pre-BCR+ leukemia and accordingly, showed a much smaller cell cluster matching p53 regulated gene sets (Figure S4E, scRNAseq profile of AZD1775 treated Nalm-6 cells). In addition, the activation of caspase 3/7 was higher in KMT2A-r cells compared to non-KMT2A-r cells (Figure S6E, left). Since p53 also modulates the senescence program in response to replication stress, we also quantified senescence following AZD1775 treatment. Indeed, we found an increased proportion of SA-β-Gal-positive cells in AZD1775 treated KMT2A-r compared to non-KMT2A-r cell lines (Figure S6F, right), congruent with increased expression of double-stranded DNA (dsDNA) sensor pathway (*cGAS*, *STING*) and components of the senescence-associated secretory pathway (SASP, *HMGB2*) in RS4;11 cells (Figure S4F-G) and high gene set score for senescence pathway (Figure S6G). Taken together, these data demonstrate a higher drug tolerance of pre-BCR+BCL6+ cells (cell fate III).

### Sequential drug treatment can target non-genetic evolution of leukemic cells

To examine the candidate cell state markers for drug tolerance (Figure 6) in primary KMT2A-r cells, we analyzed the early transcriptional changes to single-agent AZD1775 treatment in the MLL-7 PDX model. Cells were collected for scRNA-seq after 28 h treatment with 120 mg/kg AZD1775 or vehicle and sorted for alive CD45^+^CD19^+^ population (Figure S7A-B). Among the seven identified cell clusters (Figure 7A, refer also to Figure S7B), gene enrichment analysis revealed significant over-representation of genes from the BCR signaling pathway in the smallest cluster (cluster 6, Table S2). Resembling the profile found in RS4;11 cell state 9, expression of pre-BCR or BCR signaling, and pre-B TF genes was increased, while *MYC* levels declined in this cluster (Figure 7A). Previous work established that MEF2D, SREBF1 and BCL6 create a functional TF circuit in MEF2D-fusion ALL (Tsuzuki, Yasuda et al. 2020). Therefore, we included an *in vivo* treatment profile in a MEF2D-fusion ALL case for comparison (Figure 7B). The tightly coordinated expression of *MEF2D*, *BCL6* and *SREBF1* and several BCR-signaling genes (untreated diagnostic sample shown in Figure S7D) further supports that this transcriptional circuit is activated upon drug-induced stress and that the tolerant cell phenotype is present *in vivo* in primary KMT2A-r and in MEF2D-fusion patient ALL cells (Figure 7A-B).

**Figure 7.**
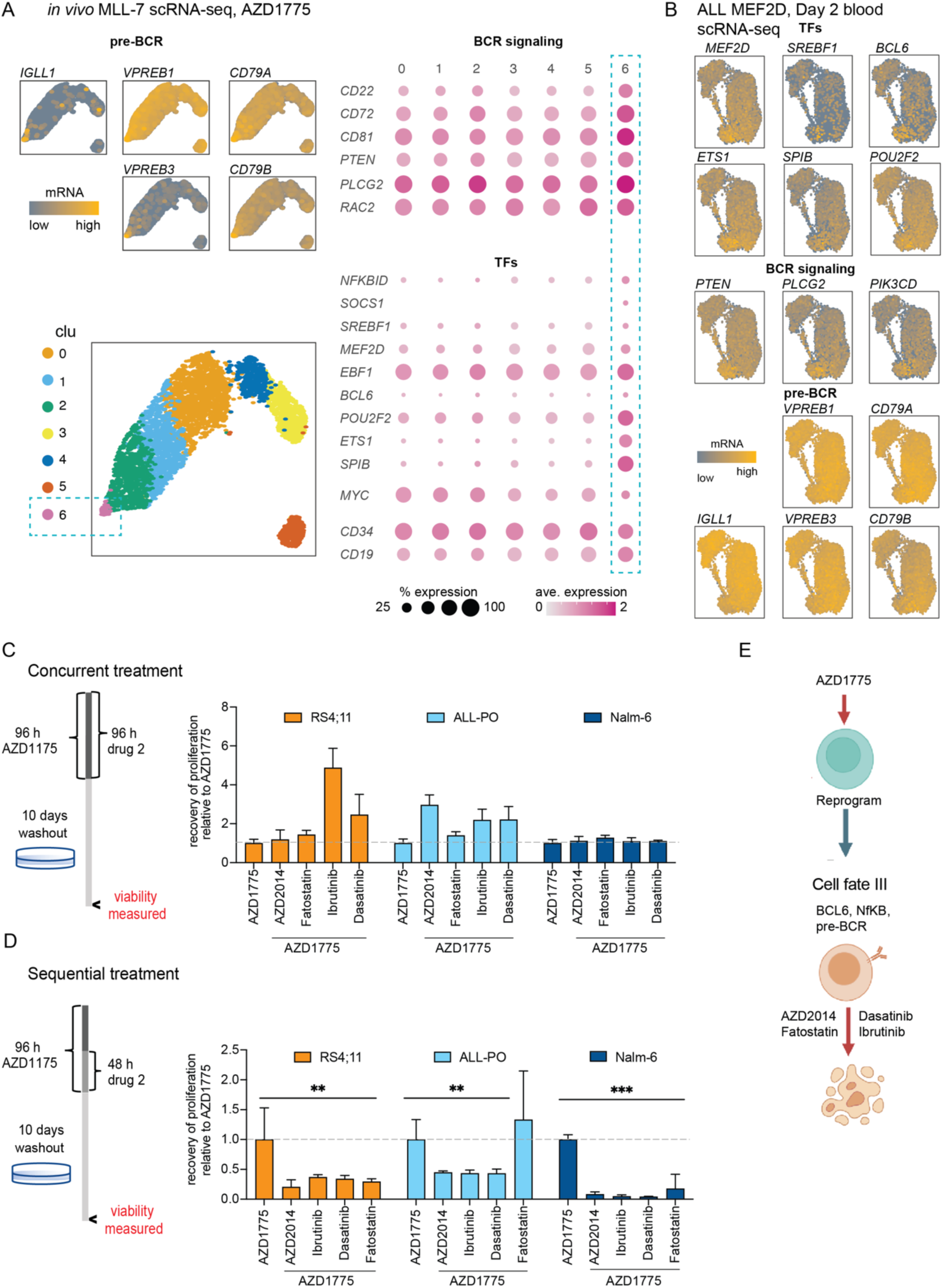
Drug tolerance-related gene regulatory circuit activation *in vivo* informs sequential drug treatment strategy. **A.** scRNA-seq transcriptome-based clustering for the primary KMT2A-r MLL-7 cells treated with AZD1775 (24 h + 4 h) is shown on the UMAP visualization. Left panel: mRNA level for pre-BCR genes (top, brighter color tones correspond to higher level) and cluster assignment (bottom). Right: dot plot heatmap showing expression levels of BCR-signaling pathway related genes and pre-B cell fate TF genes. The percentage of cells expressing each gene is indicated by dot size and average expression level by tones of red. **B.** scRNA-seq transcriptome-based clustering for primary MEF2D-fusion ALL cells at day 1 after treatment start (standard chemotherapy). mRNA level is visualized on the UMAP as in 7A. **C.** Left panel: Schematic of concurrent drug treatment experiment. Cells were treated for four days with AZD1775 in combination with the indicated drugs and allowed to recover in drug-free media for an additional 10 days. Right panel: recovery of cell proliferation after concurrent treatment, following removal of AZD1775. Drug response with indicated drugs relative to recovery of proliferation relative to AZD1775 mono-treatment is shown. Regrowth was assessed using Alamar Blue stainings. **D.** Left panel: Schematic of sequential drug treatment experiment. Cells were treated for two days with AZD1775 following by co-culturing with indicated drugs for two days for 48 h and allowed to recover for an additional 10 days without the drugs. Right panel: recovery of cell proliferation after sequential treatment, following removal of AZD1775. Drug response relative to recovery of proliferation relative to AZD1775 mono-treatment is shown (See fig S1 C). Regrowth was assessed using Alamar Blue stainings. ** denotes p<0.01 and *** denotes p<0.001 between AZD1775 alone and indicated drug combinations determined using Student’s t-test. Data are represented as mean ± SD. **E.** Schematic model of the treatment strategy targeting the regulatory circuits of induced drug tolerance cell state in response to WEE1 inhibition, Created with BioRender.com

Drugs disrupting this regulatory network activated in cell fate III could have high efficacy in preventing recovery of the remaining leukemic cell population. Therefore, we selected drugs that target pre-BCR signaling (dasatinib and ibrutinib) (Bicocca, Chang et al. 2012, Kim, Hurtz et al. 2017) and based on higher TF access observed in cell state III for master regulators of cellular lipid metabolism such as SREBF1, included also inhibitors of metabolism (mTOR inhibitor AZD2014) and fatty acid synthesis (fatostatin) (Kamisuki, Mao et al. 2009, Basu, Dean et al. 2015). To assess the potential of repositioning these drugs as effective combination therapies with AZD1775, we first measured efficacy of each drug as single agents in a drug washout setting (illustrated in Figure 7C-D, Figure S7C) and quantified the ability of cells to recover, as this experimental setup better mirrors the ability of drug-tolerant cells to recover post-treatment. As monotherapies, these drugs did not affect recovery of proliferation, and interestingly (Figure S7C), nor when administered concurrently in combination with AZD1775 (Figure 7C). Supporting successful targeting of adaptive survival pathways, the sequential administration of AZD1775 with the tested drugs effectively prevented cell survival of KMT2A-r RS4;11 and ALL-PO cells (Figure 7D). Notably, administration of AZD1775 followed by AZD2014, dasatinib or ibrutinib also strongly attenuated recovery of non-KMT2A-r Nalm-6 cells (Figure 7D).

Taken together, these results show that perturbing cell cycle regulation through inhibition of WEE1 kinase in KMT2A-r cells triggers multiple distinct genome-wide responses that include rewiring of TF programs, resulting in cell state diversification. Sequential administration of AZD1775 with drugs targeting metabolism and pre-BCR signaling can disrupt the non-genetic evolution to drug tolerant state, thus providing new opportunities for drug repositioning in preventing leukemic cell recovery (Figure 7E).

## Discussion

Targeted therapy holds promise in cancer treatment to reduce the toxicity of long-term chemotherapy. However, the acquisition of cancer cell phenotypic changes driven by tolerance pathways and emergence of acquired resistance is a common drug escape mechanism that occurs in various cancer types in response to a range of targeted treatments. In the present study, we characterized in detail the genome-wide response and cell state evolution upon cell cycle checkpoint-targeted therapy through inhibition of WEE1 kinase in ALL. Data-driven analysis from the HEMAP and the DepMap genomics resources combined with drug response profiling in different ALL subtypes revealed higher expression and increased WEE1 gene dependency in KMT2A-r compared to other subtypes. We show that WEE1 inhibition and CDK1 activation led to degradation of RUNX1 and that protein levels of other key TFs regulating the KMT2A-r leukemic phenotype were decreased. By comprehensively profiling the drug response dynamics using primary transcription and single cell transcriptome and chromatin readouts at different time points, we demonstrate that a more drug tolerant cell state was induced in surviving cells through a TF network upstream pre-BCR and BCR signaling genes. We confirmed that this drug-tolerant cell state was present in a primary KMT2A-r patient-derived xenograft model and resembled a TF circuit found in MEF2D-fusion positive ALL. The systematic characterization of the drug response at the regulatory network level afforded an opportunity to design specific drug combinations that when administered sequentially with WEE1 inhibitor prevented recovery in ALL cells.

Upon WEE1 inhibition, most of ALL cell lines, including KMT2A-r cells, rapidly enriched for cell cycle markers consistent with deregulated DNA replication and unscheduled transition into G2M-phase, leading to cell death. However, a diversification of cell states occurred at transcriptional and epigenetic level, already after 10 h treatment. Three distinct cell fates emerged in response to AZD1775 exposure in RS4;11 cells: I) extended mitotic arrest distinguished by condensed chromatin status, II) strong p53 activation leading to induction of apoptosis and senescence pathways and importantly, III) a drug tolerant cell state with higher chromatin access at TF motifs of NfKB, MEF2D and the metabolic regulators SREBF and LXR, a TF circuit regulating highly correlated expression of BCL6 and pre-BCR genes. Based on the single cell assays and primary transcription activity, this drug-tolerant cell state was induced upon WEE1 inhibition.

The comparison of enhancer activation signals in the cell fate-specific open chromatin at p53, RUNX1 and NFkB bound sites provided insight into how the alternative cell fates could arise downstream TF activity changes: a progressive response activating p53-bound sites in cell fate II diverted cells with unscheduled S-phase entry away from cell cycle and towards cell death and senescence through p53 activation at initially transcriptionally silent regions. As revealed by pulse-chase labeling of replicating cells, AZD1775 also promoted premature entry into mitosis from S-phase. The condensed chromatin state (cell fate I), also visible in scATAC-seq, remained unresolved still at 32 h. The delayed mitotic exit and replication stress response activating p53 are consistent with previously recognized drug mechanism of action (Elbaek, Petrosius et al. 2020) (Moiseeva, Qian et al. 2019). However, loss of chromatin access at RUNX1 occupied sites and activation of cell fate III-specific chromatin co-localizing with NFkB sites represent a previously unrecognized connection between cell cycle and cell fate regulatory circuits.

Remarkably, the p53-driven response anti-correlated with TFs that participate in the core KMT2A-r gene regulatory network, including RUNX1, GATA2 and MYC (Cai, Gao et al. 2015, Harman, Thorne et al. 2021). Both MYC and RUNX1 are crucial drivers of leukemogenesis and regulatory targets of KMT2A-r fusions (Guenther, Lawton et al. 2008, Wilkinson, Ballabio et al. 2013) and in particular KMT2A-r cells display strong addiction to RUNX1 for survival (Wilkinson, Ballabio et al. 2013, Wray, Deltcheva et al. 2022). The loss of RUNX1 through protein degradation may impinge upon multiple surveillance mechanisms that when disrupted enhance the sensitivity of KMT2A-r cells to AZD1775. RUNX proteins have been reported to function directly as integral regulators of DNA repair via the Fanconi Anemia pathway (Tay, Krishnan et al. 2018), while previous studies demonstrated that KMT2A-r cells have compromised the ATR-mediated S/G2-phase checkpoint (Liu, Takeda et al. 2010, Liu, Takeda et al. 2010). Consequently, inhibition of WEE1 in KTM2A-r cells may not only override the G2/M-phase checkpoint but also cause disruption of the G1/S checkpoint (Moiseeva, Qian et al. 2019), resulting in CDK1/2-driven unscheduled origin firing, p53 activation and senescence/apoptosis. Accordingly, the critical function of WEE1 kinase in cell cycle/checkpoint regulation and increased dependency on RUNX1 in KMT2A-r is underscored by the predominant cell fates associated with apoptotic or senescent states in this subtype.

In this study, we used the genomics profiles to examine non-genetic mechanisms underlying drug tolerance. Cells capable of counteracting excessive replication stress and p53-driven apoptosis could be present or acquire tolerance via rewiring of their intrinsic TF-network. Our results indicate that WEE1 inhibition critically impacted the ability of KMT2A-r cells to sustain the RUNX1-GATA2-MYC driven regulatory program. Upon disruption of this progenitor program, alternative TFs could reroute cell fate decision favoring survival in absence of RUNX1-MYC. Notably, we found increased chromatin access at several TF motifs typically activated at later differentiation stages (Mehtonen, Teppo et al. 2020) and correspondingly the expression and accessibility of genes encoding the pro-survival BCR-and PI3K signaling components, including co-expression of BCL6 and pre-BCR signaling. Although KMT2A-r leukemias resemble the early lymphoid/multipotent progenitors by immunophenotype, recent single cell characterization of patient cells has revealed that the transcriptional programs of a subset of cells present in primary bone marrow tissue match more mature pre-B-like differentiation states (Chen, Yu et al. 2022, Khabirova, Jardine et al. 2022). Moreover, flow-sorted sub-populations of blasts at different stages of immunophenotypic maturation and cellular barcoding studies have established equipotent capacity to propagate leukemia (le Viseur, Hotfilder et al. 2008, Rehe, Wilson et al. 2013, Elder, Bomken et al. 2017). Similarly, our previous single cell genomics study deciphering features of drug resistance directly in ALL patients during induction chemotherapy (Mehtonen, Teppo et al. 2020) demonstrated that although the diagnostic blasts (ETV6-RUNX1 subtype) matched pro-B-like cell state, the persisting leukemic cells at day 15 are re-programmed towards pre-B TF activity state, highlighting the clinical relevance of a non-genetic route to drug tolerant state.

The TF activity changes that characterized the more drug-tolerant cell state III included increased NfKB TF motif access at chromatin and elevated BCL6 expression, both corresponding to TFs known to counteract p53-driven apoptosis and repress genes involved in sensing or responding to DNA damage (Webster and Perkins 1999, Phan and Dalla-Favera 2004, Ranuncolo, Polo et al. 2008). BCL6 and pre-BCR-signaling can form an oncogenic feedback loop in ALL cells that promotes survival signaling via SRC family kinases, SYK, ZAP70, and downstream PI3K activation (Geng, Hurtz et al. 2015). High BCL6 at diagnosis was proposed as a prognostic biomarker (Hurtz, Chan et al. 2019). However, we show that upon genotoxic stress, cells could rapidly switch to BCL6+ state, indicating that BCL6 might prime cells for the emergence of acquired resistance during continuous treatments, and thus represent a dynamic biomarker that could be monitored at early treatment time points. Due to the recent development of efficient drugs inhibiting downstream pathways (dasatinib, ibrutinib), this elevated drug tolerance could potentially be disrupted. Our results further implicate regulation of lipid metabolism (LXR, SREBF) as a concomitant change that has been recognized in immune cell activation and oncogenic PI3K signaling in different cancers (Bensinger, Bradley et al. 2008, Ricoult, Yecies et al. 2016). MEF2D-fusions that define a rare ALL subtype with unfavorable prognosis harbor constitutive activity of a TF circuit including BCL6 and SREBF1 (Tsuzuki, Yasuda et al. 2020). Here, we compared the *in vivo* drug response in a primary KMT2A-r xenograft and a MEF2D-fusion patient sample. The highly correlated gene expression of the respective TFs and their target genes gives prominence to the relevance of this pro-survival circuit as a tolerance mechanism upon cell cycle targeting (AZD1775, or vincristine, respectively). By interfering with pro-survival signaling through targeting lipid metabolism or inhibition of BCR signaling, sequential administration of mTOR inhibitor AZD2014, fatostatin and even more potently inhibitors of BCR-signaling (dasatinib, ibrutinib), completely blocked recovery of KMT2A-r *in vitro*. Notably, this strategy strongly reduced recovery also in Nalm-6 cells, indicating that sequential targeting of WEE1 and BCR or metabolic activity, may be effective more broadly in pre-BCR+ ALLs.

Our results, consistent with a number of ALL drug therapy studies, indicate that targeting specific cancer vulnerabilities using carefully designed drug scheduling, may represent the most useful measures for improving leukemia treatment success and counteracting disease recurrence. Despite existing data of potent chemo-sensitizing activities of AZD1775 in combination with different cytotoxic agents (Tibes, Bogenberger et al. 2012, Van Linden, Baturin et al. 2013, Ford, Baturin et al. 2015, Garcia, Snedeker et al. 2017), WEE1 inhibitors have not been tested, to our knowledge, in ALL clinical trials. To date, several ongoing efforts aim to introduce immunotherapy in B-ALL treatment which requires implementation of a successful chemotherapy regime to initially reduce the leukemia burden, followed by immunotherapy. Combination therapies with WEE1 inhibitors may be attractive in this setting, reducing leukemia burden by inducing cell death through the cell-intrinsic mechanism, as characterized here. In solid cancers, favorable responses have been reported in phase 3 trials, though concerns in toxicity remain when combined to strongly cytotoxic backbone drug therapies. Our results show that several low toxicity drugs, already in clinical use for ALL or other non-communicable disease, potentiate the efficacy of AZD1775 in leukemia to overcome this challenge. Furthermore, AZD1775 was recently reported to additionally activate immune signaling through activation of STING and STAT1 pathways, or the double-stranded RNA viral defense pathway, and sensitize solid tumors to immune checkpoint therapy (Guo, Xiao et al. 2022, Taniguchi, Caeser et al. 2022). Pre-clinical testing in xenograft models lacking the immune system, as used here, may therefore only partially represent the efficacy of AZD1775 monotherapy. In this study, we demonstrated that drug scheduling has a critical impact on treatment efficacy. Concurrent drug combinations with WEE1 inhibition were unable to counteract cell recovery. This result supports a model where the drug-induced persister cell state can be eradicated only succeeding activation of the cell-intrinsic signaling pathways founding a drug-tolerant cell state. Relating drug responses to functional cell fate mapping will therefore be critical in the future, and precision treatment could be implemented as first line therapy based on anticipated adaptive cellular response dynamics (Sicklick, Kato et al. 2021, Alkhatib, Rubinstein et al. 2022).

In summary, the combination of biochemical assays and cellular resolution genomics analyses provided new insight on pronounced remodelling of the gene regulatory landscape upon cell cycle checkpoint targeting. Based on the results, we propose a strategy for WEE1 inhibition in ALL that anticipates the TF network re-wiring and switch to a pre-BCR+BCL6+ cell state and targets the drug-tolerant phenotype through novel sequential drug administration, establishing a proof-of-concept that could be utilized in the design of new targeted drug screens and clinical trials.

## METHODS

### Cell culture

A panel of ALL cell lines consisting of T-ALL: T-ALL1, Peer, Molt-16 and B-ALL cell lines: Nalm-6 (ETV6-PDGFRB); 697, RCH-ACV, Kasumi-2, MHH-Call3 (TCF3-PBX1); REH (ETV6/RUNX1); RS4;11, SEM, ALL-PO, KOPN8 (MLLr-ALL); SupB15, TMD5 (BCR-ABL) were purchased from the The Department of Human and Animal Cell Lines (DSMZ). Cell lines were maintained in RPMI1640 (Gibco, Thermo Fisher) supplemented with 10% FBS (Gibco, Thermo Fisher) and 2 mM L-glutamine (Gibco, Thermo Fisher) and 10 mM HEPES (Gibco, Thermo Fisher).

### Drug and dose response assessment

AZD1775, AZD2014 (Astra Zeneca), Dasatinib, Ibrutinib and Fatostatin (MedChemExpress, Europe), were reconstituted in dimethyl sulfoxide. Drugs were stored in aliquots at −20 °C. Alamar Blue assay was used to determine cell viability of the cells treated with increasing doses of drugs for 72 hours. Cells (30000 cells/well) were seeded into 96-well plates. Following drug treatment as indicated for 72 h, cell viability was measured using a Alamar Blue assay according to the manufacturer protocol. For the recovery of proliferation cells were treated for 72 or 96 hours and allowed to recover for an additional 10-14 days without the drugs. Regrowth was assessed by Alamar Blue staining. Absorbance was measured at 760 nm by spectrophotometer. All experiments were repeated three times.

**Analysis of apoptosis using fluorescence live cell microscopy** Cells were stained with Incucyte® Caspase-3/7 Dye for Apoptosis (Sartorius) according to the manufacturer’s instructions, followed by live cell microscopy performed with an IncuCyte S3 Live Cell Analysis System (Essen Bioscience). Nine planes of view were collected per well, using the 20x objective. The obtained data were analyzed with the IncuCyte S3 Cell-by-Cell Analysis Software Module (Essen Bioscience).

### Western blotting

For biochemical analyses of protein phosphorylation and total level, the cells were lysed in NP-40 lysis buffer (50 mM Tris-HCl pH 8.0, 150 mM NaCl, 1% NP-40) supplemented with both protease (Complete mini, Roche) and phosphatase (PhosSTOP, Roche) inhibitors. Proteins were resolved in-12% Bisacrylamide gel under reducing-denaturing condition, blotted on to PVDF membrane, and detected by immunoblotting using the appropriate antibodies. The list of antibodies and their sources can be found in the appendix file.

**Analysis of bulk gene expression data in HEMAP**. Normalized and log-transformed gene expression levels were compared between different hematologic malignancies or ALL subtypes based on the Wilcox test (Polonen, Mehtonen et al. 2019).

### PDX transplantation and AZD1775 treatment

In-house bred NOD.Cg-*Prkdc^scid^ Il2rg^tm1Wjl^*/SzJ (NSG) mice of 8-9 weeks were sublethally irradiated (250cGy) and 24 hours later, mice were transplanted through the tail vein with 2·10^6^ cells from the MLL-7 donor, established from a diagnostic sample from a child with ALL and the *KMT2A::AFF1* fusion gene (courtesy of Dr. Richard Lock) (Lock, Liem et al. 2002) Ciprofloxacin (KRKA, Stockholm, Sweden) was given in the drinking water for the duration of the experiment. A 1% engraftment of hCD45^+^hCD19^+^ cells was set as a starting point for the treatment. Engraftment was checked by taking 60 µl of blood from *vena saphena* 14 days after transplantation and then engraftment was reassessed every other day if the 1% hCD45^+^hCD19^+^ was not reached. Red blood cells were lysed with ammonium chloride (Stemcell technologies, Cambridge, United Kingdom), washed in PBS (GE Healthcare Life Sciences, Logan, UT, USA) + 2% FBS (Thermo Scientific, Waltham, MA, USA), blocked with 10% mouse serum (#M5905, Sigma Aldrich) and stained with the following antibodies: anti-human CD45-APC (clone HI30, #555485, BD Bioscience, Franklin Lakes, NJ, USA) (hCD45) and anti-human CD19-APC/Cy7 (clone HIB19, #302217, Biolegend, San Diego, CA, USA) (hCD19). Dead cells were excluded with Draq7 (Biolegend). Flow cytometric analysis was performed using the LSRFortessa (BD Biosciences) and data analysis using the FlowJo software (FlowJo, LLC, Ashland, OR, USA). Once the desired engraftment of 1% of human cells was achieved, mice (N=3 per group) were either treated with 120 mg/kg of AZD1775 (#HY-10993 MedChemExpress, Monmouth Junction, NJ, USA) 5 days a week, for 21 days (5 days on, 2 days off) (Figure S1D-H) for the survival experiment, or 48 hours with 120 mg/kg AZD1775 (#HY-10993) every 24h, and 4 hours after the last dose. Before the mice were sacrificed, blood was taken from *vena saphena*, and mice were then euthanized by cervical dislocation. Upon sacrifice, an autopsy was performed and bone marrow cells from the tibia, femur, and hip from both legs, as well as from the spleen, were collected and homogenized into single-cell suspensions by manual trituration and viably frozen in 10% DMSO (Sigma-Aldrich, Saint Louis, Missouri, USA) in FBS (Thermo Scientific). Mice were bred and maintained in accordance with Lund University’s ethical regulations and approved by the local ethics committee of Lund, Sweden.

**GRO-seq assay.** To analyze the actively transcribing RNA polymerases genome-wide, GRO-seq assays were performed as in (Heinäniemi, Vuorenmaa et al. 2016). Five million nuclei were collected from RS4;11 cells after 4, 10 and 24 h AZD1775 or DMSO treatment. The nuclear run-on reaction buffer (496 mM KCl, 16.5 mM Tris-HCl, 8.25 mM MgCl2 and 1.65% Sarkosyl (Sigma-Aldrich, Steinheim, Germany) was supplemented with 1.5 mM DTT, 750 μM ATP, 750 μM GTP, 4.5 μM CTP, 750 μM Br-UTP (Santa Cruz sc-214314A, Biotechnology, Inc., Dallas, Texas, USA and Sigma-Aldrich B7166) and RNAse inhibitors (RNase Inhibitor (Thermo Fisher, Carlsbad, CA, USA, and RNasin® Plus RNase Inhibitor Promega). To each 100 μl of nuclei samples 50 μl of the run-on reaction buffer was added and incubated for 5 min at 30°C. The RNA was collected using Trizol LS, fragmented and run-on reaction products were immuno-purified two times using anti-Br-UTP antibody (ab-6326, Abcam) bound to 30 μl of magnetic beads (Protein G DynabeadsThermo Fisher Scientific) and washed with 300 μl of PBST wash buffer four times (refer to detailed protocol in Roberts et al. 2015). The cDNA template was PCR amplified (Illumina barcoding) for 12 cycles and size selected to 225–350 bp length. The ready libraries were sequenced with Illumina Hi-Seq2000 (GeneCore, EMBL Heidelberg, Germany).

**ChIP-seq.** To distinguish the signal corresponding to active enhancer regions from the collected GRO-seq profiles, RS4;11 cells were similarly treated for 4, 10 and 24 h with AZD1775 or DMSO and cells were crosslinked with 1/10 of volume of 11% formaldehyde solution (37% formaldehyde (Sigma-Aldrich), 5M NaCl, 0.5 M EDTA pH 8.0, 0.5M EGTA, 1M HEPES) for 10 mins gently rotating at 2 mio/ml cell concentration in media. The reactions were quenched by adding glycine to a final concentration of 120 mM, and cells were washed twice with ice-cold PBS. Crosslinked lysate was flash frozen with liquid nitrogen and stored at-80°C. Lysates were thawed, nuclei were extracted, and MNase treated and antibody bound chromatin collected by adding 25 μl of Dynabeads Protein G (Thermo Fisher Scientific) and rotating for 1 hour at 4°C as previously described (Heinäniemi, Vuorenmaa et al. 2016) with the following minor modifications: 5 U-1U of MNase (Thermo Fisher Scientific) was used and 5 cycles sonicated then (Bioruptor® Plus and Bioruptor® Next Gen, Diagenode). Then supernatant was diluted to dilution buffer 2.5 times. Samples were pre-cleared with Protein G Dynabeads (Thermo Fisher Scientific) using 25μl per sample (1 h 4°C rotating). Subsequently, input samples were taken. 5 μg of the specific RUNX1 antibody (cat# ab23980, Abcam) for 5-16 million cells and 2.5 μg of specific H3K27ac antibody (cat# ab4729, Abcam) for 5-9 million cells were used. Beads were washed with 1 ml of cold wash buffers wash buffers I, II and III as in for 3 minutes (Heinäniemi, Vuorenmaa et al. 2016). Finally, beads were washed twice with TE-buffer. Beads were eluted twice in elution buffer (1% SDS, 100mM NaHCO3). Immunoprecipitated chromatin was reversecrosslinked by adding NaCl for a final concentration of 0.2M and 0,4mg/ml of RNAse A (Invitrogen) was added to each sample for 16h at 65°C. Proteinase K treated by adding 31μl of Proteinase K buffer (161mM EDTA, 645mM Tris pH7.4, Proteinase K 0,65mg/ml (Thermo Fisher Scientific)) for 1 h at 50°C. For DNA purification, ChIP DNA Clean & Concentrator (ZymoResearch, Irvine, CA, USA) was used like manufacturés protocol. ChIP-Seq libraries were prepared as previously described (Heinz, Benner et al. 2010, Heinäniemi, Vuorenmaa et al. 2016) with samples PCR amplified for 13-14 cycles and size selected for 225-350 bp fragments by gel extraction. Single-end sequencing (75 bp) was performed with Illumina NextSeq 500/550 High Output Kit v2.5.

**GRO-seq read processing** GRO-Seq reads were quality controlled (FastQC). Reads were adapter trimmed using HOMER (version 4.9 (Heinz, Benner et al. 2010) and filtered (min 95% of positions have a min phred quality score of 10) using the FastX toolkit (http://hannonlab.cshl.edu/fastx_toolkit/). Reads mapping to rRNA regions (AbundantSequences as annotated by iGenomes) were discarded. The Bowtie software (version 1.1.2 (Langmead, Trapnell et al. 2009)) was then used for alignment of remaining reads to the hg19 genome version, allowing up to two mismatches and no more than three matching locations. The best alignment was reported. Reads corresponding with so-called blacklisted regions that include unusual low or high mappability as defined by ENCODE, ribosomal and small nucleolar RNA (snoRNA) loci from ENCODE and a custom collection of unusually high signal depth regions from GRO-seq was used to filter the data. GRO-seq tagDirectories were generated from aligned reads with fragment length set to 75 and data visualized using makeMultiWigHub.pl with strand-specificity HOMER (version 4.9).

**ChIP-seq read processing** Reads with poor quality bases scores were filtered (min 97% of positions were required to have a min phred quality score of 10) using the FastX toolkit (http://hannonlab.cshl.edu/fastx_toolkit/). Duplicate reads were collapsed using fastx (collapse). Bowtie (version 1.2.3 (Langmead, Schatz et al. 2009)) was used to align the reads to the hg19 genome version, allowing up to two mismatches and no more than three matching locations. tagDirectories were generated with HOMER (version 4.9) tool. Histone peaks (GSE148195) were identified using HOMER (version 4.9) findPeaks with setting-style histone. The peak signal profile across pooled tagdirs was then used to distinguish dips that correspond to nucleosome free regions that can be accessed by TFs (this was found to work well for peaks < 7.5 kb). To identify RUNX1 peaks, HOMER (version 4.9) findPeaks-style factor was used with the following less stringent settings: fold enrichment over input tag count 2, poisson p-value threshold relative to input tag count 0.001, fold enrichment over local tag count 2, Poisson p-value threshold relative to local tag count 0.001, false discovery rate 0.01,-tagThreshold 10. GM12878 NfKB (Tnfa treatment) from ENCODE SYDH ChIP-seq tagDirectory was generated from bam file using HOMER.

**GRO-seq gene signal analysis** To compare the effect of AZD1775 on primary transcript levels across the collected time points Refseq (genome_annot_hg19 refGene_2018) coding and non-coding gene coordinates were quantified without exon regions. A raw count matrix was generated using Homer (version 4.9.1) specifying as maximum read per position 3. not allowing combining quantified reads if the same transcripts is detected more than one time. The low expression transcripts were filtered based on cpm and rpkm values requiring a row sum greater than 0.5 in at least two samples. To detect differentially expressed transcripts, a generalize linear model was fit using functions available in the R/Bioconductor package edgeR. Genes with significant change between any of the conditions (adjusted p-value < 0.001) were reported based on one-way ANOVA-like the likelihood ratio tested with glmLRT function. The genes were then clustered using k-means (into six clusters) and plotted as a heatmap (ComplexHeatmap R package) with six clusters. One cluster profile was excluded as it reflected mainly changes in DMSO basal levels between time points. Pathway analysis was performed on the gene clusters with > 100 genes, otherwise the genes were plotted and examined individually.

**GRO-seq enhancer signal analysis** The nascent transcriptome result allows quantification of eRNA signal that can guide the analysis of TF activity. Enhancer regions are typically defined based on multiple genomic signal features. Here we started with DNA hypersensitive sites (DHS from Encode Duke narrow peaks, available across the ENCODE cell collection) and ATAC-seq peak coordinates (from RS4;11 cells in GSE117865 and from bone marrow cell populations in (Ulirsch, Lareau et al. 2019)) for specifying candidate enhancer centers. The candidate regions were extended +/-500 bp and the 1 kb size enhancer regions were quantified with Homer (homer/4.9.1) analyzeRNA. Coordinates with overlap with gene sense-transcription or the promoter region signal were excluded. Enhancers that passed a minimum cpm cutoff or intersected (bedtools2/2.27.1) with RS4;11 histone ChIP-seq peaks for H3K4me1 (GSE71616) and H3K27ac (our own GSE148195, GSE71616 and GSE117865) were kept for analysis. Intergenic enhancers should have bidirectional and approximately equal signal from plus and minus strand. If the mean difference was over 10-fold only the lower signal was considered (the stronger signal typically derives from transcribed gene region signal extending beyond annotated transcription termination site). For tightly clustered enhancers, the enhancer that overlapped best with RS4;11 ATAC or centered nuclear free region (< 7.5kb peak size) of H3K27ac peaks was selected, or otherwise the region with maximum signal was used. Quantified enhancers were filtered (expression cut-off 5 in minimum two samples) and normalization using RLE. A quasi-likelihood negative binomial generalized log-linear model was fitted using edgeR. The quasi-likelihood (QL) F-test was used to detect significant eRNA level changes (FDR < 0.1) Clustering was performed using hierarchical clustering and distinct cluster profiles were visualized using the R package ComplexHeatmap. Denovo motif discovery was performed by extracting the DNA sequence for motif enrichment analysis with Homer (4.9.1) findMotifsGenome.pl (region size 200bp, repeat masked genome, background all enhancers from statistical analysis. To further visualize enhancer signal GRO-seq signal histograms were generated. For TF binding sites, intergenic region corresponding to motif-centered TF ChIP-seq peaks were used for signal summary. p53 ChIP peaks were retrieved from GSE46991 and centered with TP53 motif (p63(p53)/Keratinocyte-p63-ChIP-Seq(GSE17611)/Homer) The histogram was generated with bin size 25 bp +/-2000bp from the ChIP-peak center. For chromatin state-specific enhancer analysis regions that had high/low accessibility in each chromatin state were detected using the Seurat Findmarkers function, with a minimum fold change difference of 0.1 and adjusted p-value a 0.1 To further visualize top 200 chromatin state specific enhancer signals the assay-specific signal matrixes were generated with annotatePeaks.pl tool of the HOMER package (version 4.11)(Heinz, Benner et al. 2010). The matrixes were generated with bin size 25 bp +/-1000 bp from the enhancer center. Visualization was done with image J (version 1.53t) (Schneider, Rasband et al. 2012) and average signal summary histograms generated with the same parameters visualized with base R plots (R version 4.1.0).

**scRNA-seq assay**. To analyze the drug response at cellular resolution RS4;11 or Nalm-6 cells were treated for 24 h with 1 µM AZD1775 or DMSO. Cell viability was checked using Trypan blue with Cellometer Mini Automated Cell Counter (Nexcelom Bioscience) and only viable cells were processed further. To deplete dead cells, the Dead Cell Removal Kit (#130-090-101, MACS miltenyi Biotech) was used and the column rinsed twice with 1 ml Binding Buffer to elute viable cells (97-98% viability). Single cell suspension, loading and library preparation was performed according to the ChromiumTM Chromium Single Cell 3'Reagent Kits v3 User guide CG000184 Rev A and libraries constructed using the 10X Genomics Chromium technology. Loading concentrations were 1700 cell/ul, 1000cell/ul for RS4;11 DMSO, AZD1775 cells; 500 cell/ul, 1300 cell/ul Nalm-6 DMSO, AZD1775, respectively. Each lane was loaded with 10 000 cells as pools of human and mouse cells (not part of this study) processed in parallel. Sequencing was performed in FIMM Technology Center Sequencing Laboratory, Biomedicum, Helsinki, Finland by using a NovaSeq S2 sequencer aiming 50 000 reads per cell depth.

### PDX cell staining and sorting for scRNA-sequencing

Sample preparation for scRNA-sequencing was performed by thawing viably frozen bone marrow cells, which were stained with Draq7 (BioLegend), hCD45 (BD Bioscience), and hCD19 (BioLegend), together with a PerCP/Cyanine5.5 anti-mouse CD45 (BioLegend). Approximately 20.000 alive/mCD45^-^/hCD45^+^/hCD19^+^ cells were sorted into a PBS (GE Healthcare Life Sciences) + 2% FBS (Thermo Scientific) coated tube and prepared for sequencing using the 10x Genomics platform at the Center for Translational Genomics, Lund University.

**Single cell genomics from primary MEF2-fusion ALL cells** Bone marrow or blood mononuclear cells collected at diagnosis and one day following treatment start were thawn from the viably cryopreserved samples. Cells were thawn in a 37°C water bath, immediately after adding 0.5 ml RPMI 1640 media (Thermo Fisher Scientific) + 10% FBS (Gibco)+ 20 µl DNAse (Roche 100U/µl) on top, then moving cells to 15 ml Falcon and filling the volume drop-wise, gently swirling. For samples with < 1 million cells, 10 µl of DNAse was added, filling volume to 5 ml. Cells were washed two times, then centrifuged and re-suspended in 50 µl Cell Staining Buffer (BioLegend) for counting. Subsequently, 100 000-200 000 cells per sample were processed by blocking with 5 µl of Human TruStain FcX Blocking Solution (BioLegend) for 10 min at 4°C. Antibody pool (0.25 µg per million cells per specific antibody, except 0.125 µg for CD45 and CD8; 0.1 µg per hashtag antibodies) was added and incubated for 30 min at 4°C. Samples were washed three times with cell staining buffer, counted and assessed for viability using trypan blue staining, and then sample pools were prepared, centrifuged and loaded to Chromium lanes (10x Genomics). If post-stain viability was >70% for all samples, equal ratios were loaded aiming at 10 000 cells. Otherwise, loading ratio was adjusted such that half target cell count was used for samples with viability 50-70% and one fifth of viability was 20-50%. The samples analyzed in this study (diagnosis EG9, day 2 EG19) are part of a sample set processed in three batches and with hashtag-barcoding of two donors per Chromium lane. The 5’ gene expression, ADT-and VDJ (BCR and TCR) libraries were prepared following manufacturer instructions (10x Genomics) and cDNA qualities were assessed using Bioanalyzer. Libraries were index barcoded and sequenced using Illumina Novaseq (S1/S2).

**scMultiome-seq assay** To analyze the drug response transcriptional and chromatin accessibility from the same cells, RS4;11 were treated for 10 h with 1 µM AZD1775 or DMSO. In scMultiome assay nuclei were first isolated. 10x Genomics protocol (User guide CG000365 RevB) was first optimized for the RS4;11 cell line with a 3-minute cell lysis time. Nuclei were then washed 3 times with 600 g 7 min centrifugations steps. Nuclei quality was determined with light microscoping at 40x focus. Nuclei count and lysis optimalization was determined with diluted Trypan blue solution with Countess 3 cell counter (Invitrogen). Each sample was loaded with 10 000 nuclei and library preparation performed according to the Chromium Next GEM Single Cell Multiome ATAC + Gene Expression User guide CG000338 Rev E and libraries constructed using the 10X Genomics Chromium technology. Loading concentrations were 7200 nuclei/ul for DMSO cells and 4860 nuclei/ul for AZD1775 cells. Multiome ATAC Sequencing was performed in Novogene Cambridge, United Kingdom by using a NovaSeq PE50 sequencer aiming 20 000 reads per cell depth. Multiome RNA samples were sequenced in the same place with NovaSeq PE150 aiming 50 000 reads per cell depth.

**scRNA-seq read processing** scRNA-seq data was processed and aligned with Cell Ranger (version 3.0.2) using hg19 and mm10 genome as reference. Only the human cell counts were used for future analysis performed using the R package Seurat (v3) and the Python package Scanpy (version 1.5.1). Data per cell line was filtered with the following cutoffs: cells had more than 2000 and less than 6000 UMIs, percentage of mitochondrial counts less than 20%, and less than 500 mouse mm10 gene counts. Post-filtering, RS4;11 DMSO had 4169 cells and AZD1775 sample respectively 5135 cells. Post-filtering, Nalm-6 DMSO had 5598 cells and Nalm-6 AZD1775 sample resulted in 5039 cells. Cells were log-normalized using scaling factor of 10 000.

PDX cell scRNA-seq data was processed and aligned with Cell Ranger (version 6.0.1) using GRCh38-2020-A genome as reference. Utilization of R package Seurat (v3) enabled further analysis. MLL-7 data was filtered with the following cutoffs: cells had more than 1500 but less than 6000 UMIs and percentage of mitochondrial counts less than 10%. Post-filtering, DMSO had 4432 cells and AZD1775 resulted in 4672 cells. Cells were log-normalized using a scaling factor of 10 000. Vst selection method (Seurat) was implemented to select 2000 highly variable features (genes). DMSO and AZD1775 cells were clustered using the FindNeighbors function based on PCA dimension 1:10 and with resolution 0.5. This resulted in a discovery of 7 distinct cell clusters in DMSO and treated cells.

Primary ALL scRNA-and scADT-seq data was aligned with Cell Ranger 6.0 version to human reference (hg19) with default parameters. Donor (and singlet/doublet) assignment was carried out by DSB-normalized hashtag signals and downstream analysis carried out using Seurat.

**scRNA-seq cell state analysis** Vst selection method (Seurat) was used to select 2000 highly variable features (genes). Cells were clustered using the FindNeighbors function based on PCA dimension 1:10 and with resolution 0.5. This resulted in a discovery of 11 distinct cell clusters in RS4;11 cells that are referred to as “cell states”. UMAP visualizations were used to represent the data in two dimensions. For cell cycle characterization, a list of cell cycle markers from Tirosh et al. 2015 were utilized. Several statistical comparisons between the cells assigned to these cell states were performed: i) Genes that had high/low expression in each cluster were detected using the Seurat Findmarkers function, with a minimum fold change difference of 0.25 and adjusted p-value between cells assigned to the cluster and other cells. ii) AZD1775-treatment specific cell states were compared with the cell state matched to normal cell cycle G2/M-phase (cell state 4), iii) cluster markers were detected between AZD1775-treatment specific cell states. Violin plots were used to visualize gene expression distributions. For visualizing gene expression level per cell state all unique genes from cluster comparisons were acquired and the mean scaled counts were summarized by cell type. These mean expression values were then used as input for hierarchical clustering with 1-pearson correlation as a distance metric. ComlexHeatmap R package was used for visualizations, testing different cluster numbers (10-20) for row-wise clustering. Pathway analysis was performed for the gene clusters and for broader patterns distinguished based on the heatmap.

### scMultiome-seq read processing

For comparing to 24 h data, multiome scRNA-seq data was processed with the same Cellranger preprocessing workflow, resulting in reads mapped to hg19 genome version. DMSO and AZD1775 samples were merged and analyzed using Seurat v4.1.1 using the following cutoffs in filtering: 1500 < UMIs < 6000 UMIs and percentage of mitochondrial counts < 20%, resulting in 12835 DMSO and 14788 AZD1775 cells. The raw reads were also processed and aligned with Cell Ranger Multiome workflow (cellranger-arc-2.0.1, hg38 genome version). The cell assignment available from per barcode metrics output was used to select nuclei for downstream analyses. The scATAC-seq workflow based on Signac ((Stuart, Butler et al. 2019) 0.1.6 and Seurat 3.1.1. versions that was used for 24 h data was run with minor modifications (EnsDb.Hsapiens.v86, no blacklist filtering, atac_peak_region_fragments > 1000 & atac_peak_region_fragments < 100000 & pct_reads_in_peaks > 50 & nucleosome_signal < 10) to generate comparable results. Nucleosome signal histogram analyses were performed using Signac version 1.6.0. The matching cell barcodes between scRNA-and scATAC-seq profiles from the same cells were utilized to visualize data across modalities. For example, this allowed showing the scATAC-seq nucleosome signal on the UMAP generated from scRNA-seq data. Cell type labels were assigned based on label transfer analysis with the respective 24 h sample used as reference sample.

### scRNA-seq RNA velocity and PAGA analysis

Dynamic changes in gene transcription can be modelled based on reads corresponding to both unspliced and splices mRNA (newly synthetized RNA vs processed RNA, respectively). Based on the dynamic RNA processing model (La Manno, Soldatov et al. 2018), the future transcriptome state can be visualized together with the measured current state. This analysis was used to analyze cell state dynamics in the RS4;11 cell model. Velocyto CLI (version 0.17.17 (La Manno, Soldatov et al. 2018)) was used to calculate the count matrices, masking expressed repetitive elements (available for hg19 from UCSC Genome Browser). The scVelo-package (version 0.2.1 (Bergen, Lange et al. 2020)) was used in downstream analysis. The gene expression matrix was accompanied with the spliced and unspliced count matrices. The data was first filtered by removing genes with less than 20 shared counts in both spliced and unspliced data. The matrices were each then normalized by dividing the counts in each cell with the median of total counts per cell. The 2000 most variable genes were extracted based on the spliced count matrix and the data matrices were log-transformed. Top PCs (30) were used for neighborhood graph calculation, with the number of neighbors set to 30. Then the AZD1775 and DMSO samples were separated for RNA velocity analysis. Based on the neighborhood connectivities, the first order moments for spliced and unspliced matrices were calculated and the velocity estimated using the dynamical model. The velocities were embedded to a UMAP presentation, which was calculated with 20 PCs and 10 neighbors. A PAGA graph was used to visualize connectivities (dashed lines) and transitions (solid lines). Python Scanpy-package (version 1.5.1) was used for gene set scoring. Before scoring, gene sets used in the analysis (e.g. senescence gene sets) were filtered to include only genes that are expressed in the data, excluding G2M-phase markers (to remove cell cycle bias in the scores).

**scATAC-seq assay**. RS4;11 cell nuclei were collected from parallel cell cultures as those used for the scRNAseq experiments to study chromatin accessibility at the single cell level (scATAC-seq). Nuclei were isolated with 10x Genomics Nuclei Isolation for Single Cell ATAC Sequencing user guide CG000169 Rev D. Nuclei quality was determined with light microscoping with a 40x focus. Nuclei concentration was determined with Trypan blue and pools of human and mouse (not part of this study) cell lines were loaded together. scATAC-seq was performed using Chromium Single Cell ATAC Library & Gel Bead and Chip E Single Cell ATAC Kit (10x Genomics), user guide CG000168 Rev C. Sequencing was performed in the FIMM Technology Center Sequencing Laboratory, Biomedicum, Helsinki, Finland using NovaSeq S2 sequencer aiming 50 000 reads per cell depth.

### scATAC-seq read processing

The raw reads were processed and aligned with Cell Ranger atac workflow (version 1.2.0). Seurat ((Stuart, Butler et al. 2019), version 3.1.1) package and Signac ((Stuart, Butler et al. 2019) version 0.1.6) were used for downstream analysis, selecting only the human peak coordinates and cells. Peaks detected from DMSO and AZD1775 treated cells were merged using bedtools (version 2.27.1), discarding < 10 bp wide peaks. Data across treatments was combined by re-quantifying counts from the fragment files based on the merged peak regions. Peaks detected in > 1% of cells were kept for downstream analyses. Nucleosome signal was quantified based on fragments mapping to chr 1 using the Signac package NucleosomeSignal function. TSS enrichment was calculated based on coordinate ranges retrieved from EnsDb.Hsapiens.v75. These quality metrics were used in combination with metrics to select good quality nuclei for downstream analyses (peak region fragments > 1000, peak region fragments < 100000, percentage reads in peaks > 15, blacklist ratio < 0.05, nucleosome signal < 10, TSS.enrichment > 2). To analyze the atypical nucleosome signal manifest in AZD1775-treated sample, the above filtering criteria were used without the nucleosome signal cutoff to plot the signal profiles. Term frequency-inverse document frequency (TF-IDF) normalization of peaks by accessibility was performed using the LSI implementation used by (Cusanovich, Hill et al. 2018). To reduce dimensions and perform clustering of the data matrix, singular value decomposition (SVD) was run on the TD-IDF normalized matrix, keeping 150 dimensions. Based on elbow plot analysis 50 dimensions were kept for UMAP visualization and clustering. To study more specifically chromatin activity responses at each treatment condition, separate UMAPs were created (reduction method lsi, dims=1:50, resolution 0.4). The two smallest chromatin states found in the chromatin state clustering of DMSO (c5, c6) and AZD1775 (c6, c7) treated cells differed based on quality control metrics. The smallest clusters had a low peak region fragment amount (low quality) and were discarded from downstream analyses. The clusters with elevated nucleosomal signal were reproduced in the scMultiome result and could be matched using this data to cells with a good quality RNA-seq profile. Moreover, they had a distinct TF motif activity pattern in ChromVar analysis (unlike the smallest cluster). Therefore, we kept the clusters with elevated nucleosome signal in the downstream analyses. For chromatin state-specific enhancer analysis, we started with intergenic enhancer defined from bulk genomics data and identified high/low accessibility regions in each chromatin state using the Seurat Findmarkers function, with a minimum fold change difference of 0.1 and adjusted p-value a 0.1. To further visualize top 200 chromatin state specific enhancer regions the assay-specific pseudobulk signal matrixes were generated by creating a combined tagDirectory for cells matched to each chromatin state, followed by signal quantification with annotatePeaks.pl tool of the HOMER package (version 4.11 (Heinz, Benner et al. 2010)). The matrixes were generated with bin size 25 bp +/-1000 bp from the enhancer center. Visualization was done with image J (version 1.53t (Schneider, Rasband et al. 2012)) and average signal summary histograms generated with the same parameters visualized with base R plots (R version 4.1.0).

### TF motif activity profile from scATAC-seq profile

To determine motif activity, i.e. variability in chromatin accessibility per cell, the chromVAR, ((Schep, Wu et al. 2017) version 1.6.0) tool was applied separately to each sample. The known TF motifs available in the Homer tool were scored. Subsequently, cluster specific motifs were detected using statistical analysis (Seurat FindMarkers function, log2 fold change threshold 1.5, adjusted p-value <0.0001) comparing each chromatin cluster to other clusters. Redundant motifs recognized by similar protein complexes were grouped manually and the most significant motif kept for visualization. The scaled chromatin accessibility values were used to calculate mean access per chromatin cluster and visualized using the ComplexHeatmap R package. The Signac visualization functions were used to generate genome browser track plots comparing different clusters.

### Label transfer from scRNA-seq to scMultiome-seq

To characterize similar cells between transcriptome level in two different time points the canonical correlation analysis available in the Seurat (Seurat 4.1.1) package was used. First anchors between two different time points were identified following the user guide recommendations. 24 h RS4;11 data cell states 1-11 were used as reference. Then the 24 h RS4;11 cell state labels were transferred to the 10 h multiome scRNA-seq data. Prediction scores were visualized for quality control purposes (refer to Fig. S2) and all predicted cells were applied in further analysis. The same analysis was performed to characterize two different time point chromatin levels with user guide recommendation for ATAC data. 24 h RS4;11 DMSO ATAC clusters 0-5 were used as reference for 10 h DMSO multiome scATAC-seq data. 24 h RS4;11 AZD1775 ATAC clusters 0-6 were used as reference for 10 h AZD1775 multiome scATAC-seq data. Prediction scores were visualized for quality control purposes (refer to Fig. S2G) and a prediction score < 0.3 omitted cells with a poor match from visualizations (assigned to category NA).

### Pathway enrichment analysis

Gene lists acquired bulk genome wide and scRNA-seq studies were analyzed using the online web server Enrichr (Chen, Tan et al. 2013, Kuleshov, Jones et al. 2016) for enrichment of ontology and pathway terms. The analysis was performed based on gene sets from BioPlanet 2019 and TF perturbations. Enriched terms were ranked based on lowest adjusted p-value and top terms per group visualized as dot plots (R package ggplot2).

### Cell proliferation, EdU incorporation and senescence assessment by FACS

To study the effects of AZD1775 on cell proliferation, cells were incubated with 10 μM EdU for 1 hour at 37 ⸰C to label proliferating cell population. EdU-labelling of the cells were done prior the treatment for further processing according to the kit protocol (Thermo Fisher). Following cell fixation and permeabilization, the cells were stained with pH3 (1:25, BD Bioscience), and 50 μg / mL Propidium Iodide in 1% BSA/ TBS – 0.5% Tween 20 containing 10 μg/mL RNAseA overnight at 4 ⸰C. Finally, the click-iT reaction were carried out according to the manufacturer’s protocol prior to FACS analysis. To analyze senescence, treated cells were or CellEvent Senescence Green Flow Cytometry Assay Kit (Invitrogen, C10840) and Propidium Iodide according manufacturer’s protocol.

### Quantitative real-time PCR (qRT-PCR)

To perform the qRT-PCR, RNA was prepared using RNeasy kit (Qiagen). Residual genomic DNA was eliminated from the total RNA fraction by DNase I treatment (Qiagen), according to the manufacturer’s protocol (Qiagen). 50 ng of the purified RNA was then used for cDNA synthesis (SuperScript VILO cDNA Synthesis Kit, Invitrogen, Life Technologies). Quantitative PCR were carried out using SYBR green probes as described in the appendix file (Table S5). The data were analyzed with ΔΔCt method and presented as relative expression.

### Statistical analysis

Data from biochemical assays are presented as the mean + SD from two or more independent experiments unless indicated otherwise. Statistical analyses were performed using the statistical package GraphPad Prism 8 (GraphPad Software, San Diego, CA, U.S.A. http://www.graphpad.com) using either Student’s t-test, Log-rank (Mantel-Cox), one-way analysis of variance (ANOVA) or two-way analysis of variance (ANOVA), as indicated. Genome-wide statistical analysis was performed using dedicated software (R or Python packages as indicated). Multiple testing correction was performed using Benjamini-Hochberg or Bonferroni’s post hoc test when appropriate.

### Availability of data and materials

The single cell datasets analyzed and generated in the current study will be available in Gene Expression Omnibus under the accession number when manuscript will be published. Bulk data for GRO-seq in the RS4;11 cell line will be available in GEO after published. ChIP-seq datasets analyzed and generated under GSE148195 https://www.ncbi.nlm.nih.gov/geo/query/acc.cgi?acc=GSE148195. Additional data for ChIP-seq peak analysis was downloaded from GEO GSE71616 https://www.ncbi.nlm.nih.gov/geo/query/acc.cgi?acc=GSE71616, GSE117865 https://www.ncbi.nlm.nih.gov/geo/query/acc.cgi?acc=GSE117865 and GSE46991 https://www.ncbi.nlm.nih.gov/geo/query/acc.cgi?acc=GSE46991. Bulk ATAC-seq profiles of RS4;11 cells were acquired from GEO GSE117865 and from bone marrow cell populations in (Ulirsch, Lareau et al. 2019). Upon request additional information required to reanalyze the data reported in this paper is available from the lead contact.

## ACKNOWLEDGEMENTS

The authors thank Professor Richard Lock (Children’s Cancer Institute Australia) for providing KTM2A-r PDX samples and related clinical information for this study, Kuopio and Tampere university hospital pediatric oncology clinics for prospective sample collection of pediatric leukemias during induction chemotherapy and patients consenting to participate in these studies, Professor Sui Huang (Institute for Systems Biology USA) for helpful discussions, and Janne Suhonen and Jonne Nieminen for setting up bioinformatics workflows for EASI-Genomics samples. This research was funded by The Swedish Childhood Cancer Fund, the Swedish Cancer Society, The Swedish Research Council, Karolinska Institute, Radiumhemmets Research Foundation, AstraZeneca-SLL-KI Open Innovation grant (#18122013), the Academy of Finland (321553, 310106), the European Union Horizon 2020 research and innovation programme under grant agreements No 824110 (EASI-Genomics) and ERAPERMED2018-209 (JTC2018 ERA-NET ERA PerMed), Väre Foundation, Emil Aaltonen Foundation, Cancer Foundation Finland, Jane and Aatos Erkko foundation, and Sigrid Juselius foundation. The authors wish to acknowledge CSC – IT Center for Science, Finland and UEF Bioinformatics Center, University of Eastern Finland, Finland for computational resources. The authors would like to Acknowledge Clinical Genomics Lund, SciLifeLab and Center for Translational Genomics (CTG), Lund University and <u>SNP&SEQ Technology Platform</u>, SciLifeLab Uppsala, for providing expertise and service with genomics, sequencing and analysis.

