## supplemental figures for "Sequential drug treatment targeting cell cycle and cell fate regulatory programs blocks non-genetic cancer evolution in acute lymphoblastic leukemia"

### Supplementary Figures

#### Figure S1

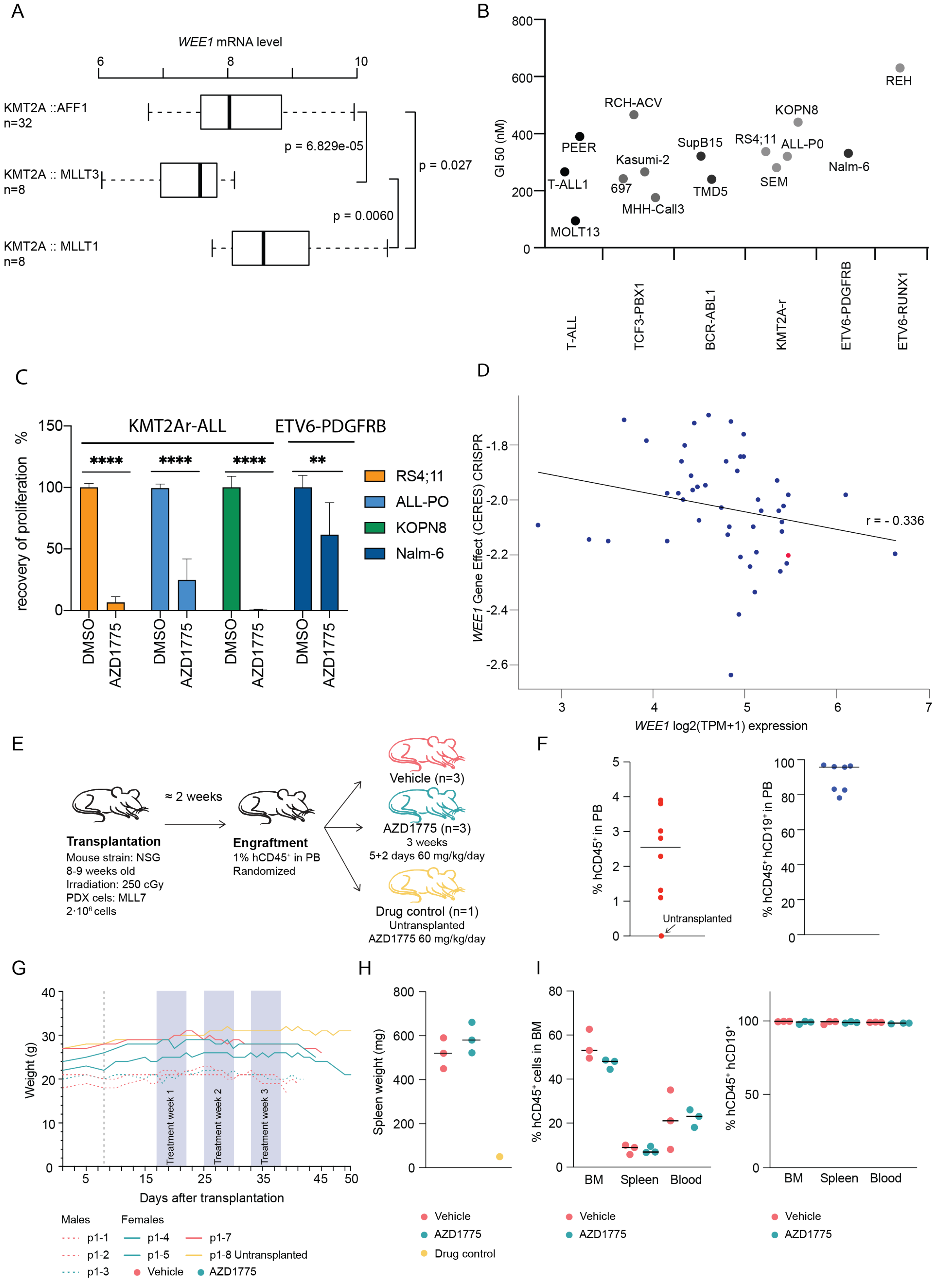


**Fig. S1, related to Fig. 1.**

**A**. Boxplots of *WEE1* gene expression level comparing different KMT2A fusions (HEMAP dataset). Wilcoxon test p-value is indicated.

**B.** GI50 values of AZD1775 treatment (72 h) comparing ALL cell lines corresponding to different genetic subtypes.

**C.** Recovery of cell proliferation following removal of AZD1775. Cells were treated for 72 hours with AZD1775 and allowed to recover for an additional 10 days without the drugs. Regrowth was assessed using Alamar Blue stainings. ** denotes $p<0.01$ and **** denotes $p<0.0001$ determined using Student's t-test. Data are represented as mean ± SD.

**D.** Scatter plot visualization of *WEE1* gene dependency based on CERES score (from DepMap) against *WEE1* mRNA level in hematopoietic cell lines. A lower CERES score indicates higher probability of dependency (0: not essential, -1 median of all common essential genes). KMT2A-r ALL cell line (SEM) is indicated in red color.

**E.** Schematic representation of the experimental plan depicting the different treatment arms.

**F.** Scatter plot showing the leukemic cell engraftment on the first day after treatment start, represented by the fraction of hCD45+CD19+ cells.

**G.** Time course of weight measurements. Mice were weighed every other day before treatment and daily afterward. Treatment was well tolerated.

**H.** Spleen weights at sacrifice (left panel)

**I.** Flow cytometric quantification of leukemia burden, showing the fraction of hCD45+ and hCD19+ cells.

Figure S2

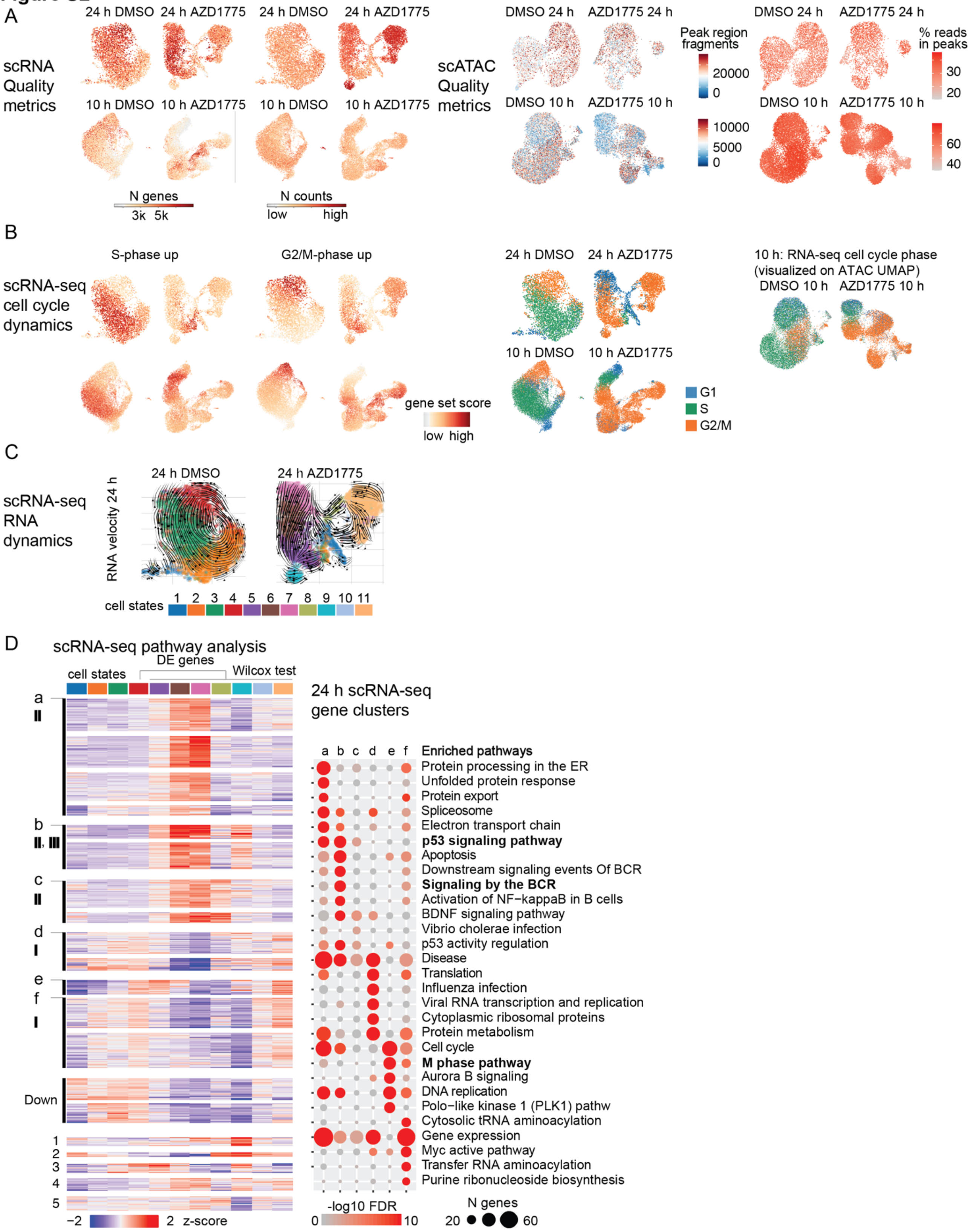


**Fig. S2, related to Fig. 2.**

**A.** scRNA-seq quality metrics (Ncounts: number of counts, Ngenes: number of genes) visualized on the RNA-seq UMAPs from DMSO and AZD1775-treated cells (top: 24 h data, bottom: 10 h data) (left). scATAC-seq quality metrics (peak region fragments, percentage of fragments in peaks) visualized on the ATAC-seq UMAPs from DMSO and AZD1775-treated cells (top: 24 h data, bottom: 10 h data) (right). Red color tones indicate higher values.

**B**. Gene set scores for S- and G2/M phase specific genes and the cell cycle state assignment based on the scores visualized on UMAPs, as in A. scRNA-seq cell cycle state labels visualized on the low dimensional UMAP representation of scATAC-seq profiles (left: DMSO, right: AZD1775).

**C**. Cell state dynamics based on RNA-velocity analysis from 24 h scRNA-seq. The colors correspond to assigned cell state based on 24 h scRNA-seq data, refer to cell states shown in Fig. 2D. Arrows point towards the predicted future state of each cell.

**D.** Differentially expressed genes from comparison of AZD1775-specific clusters (cell states 5-11) and DMSO cell state 4 (matching G2/M cell cycle phase) are shown as a heatmap where genes (in rows) are clustered based on their mean expression in cells corresponding to each cell state. The color corresponds to z-score where tones of red indicate high expression level. Pathway analysis was performed based on broader similarity of clusters (indicated by letters a-f) and shown as dot plot as in Figure 2C. Small clusters (<100 genes) with specificity to cell state 9 are indicated at the bottom. For complete gene to cluster assignment, refer to Table S2.

Figure S3

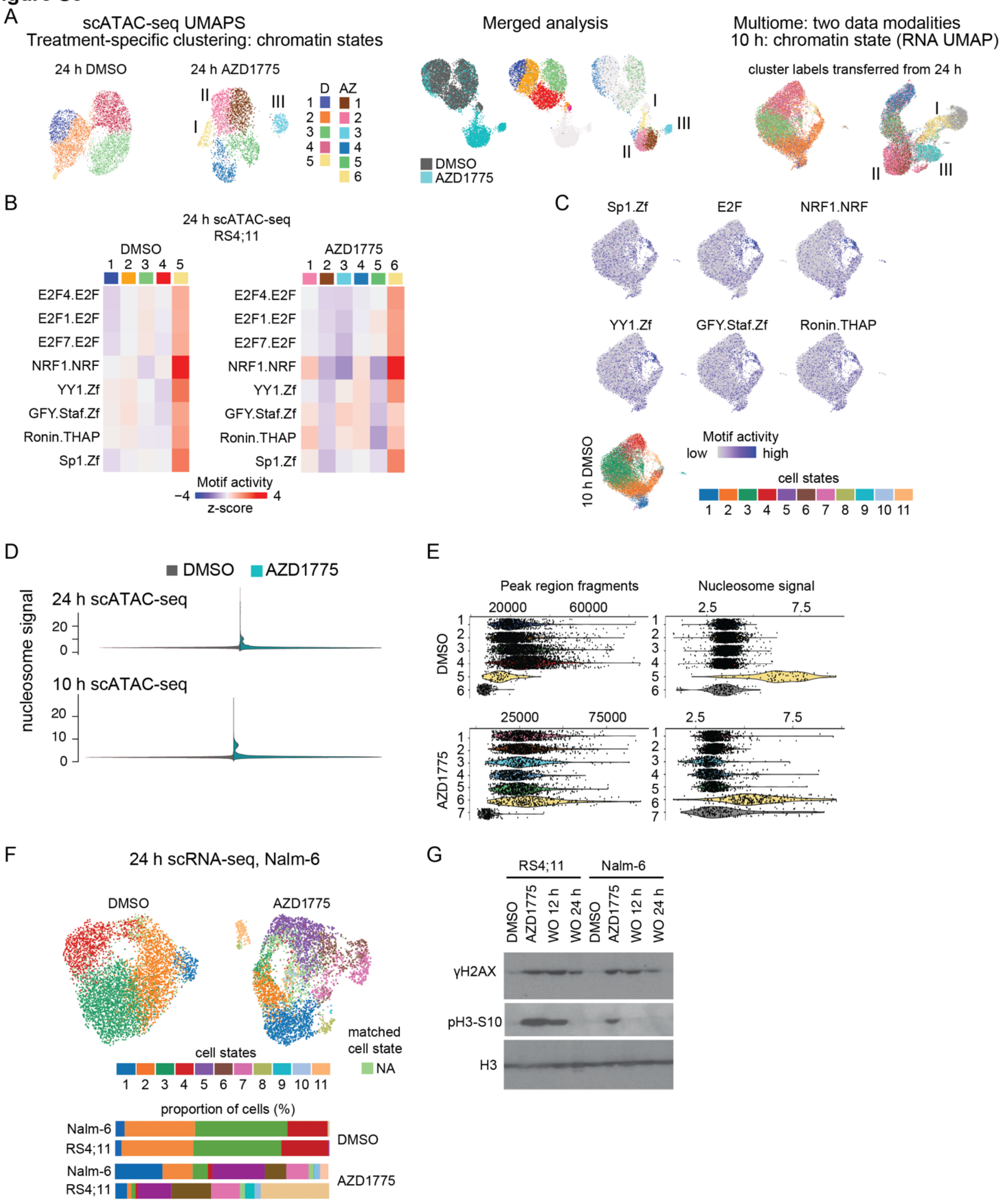


**Fig. S3. related to Fig. 3.**

**A**. Left: Low dimensional projection and scATAC-seq chromatin access-based clustering for the RS4;11 cells from DMSO and AZD1775 treated cells are shown on UMAP visualizations. Clustering of DMSO and AZD1775-treated cells using 24 h data defines chromatin states (labelled D1-5 and

AZ1-6). Middle: Merged analysis: joint representation of 24 h DMSO and AZD1775 treated cells. Right: Label transfer: cells from 10 h data assigned the label of closest 24 h ATAC profile neighbor.

**B**. Heatmap showing TF motif activity score across chromatin states in DMSO and AZD1775 treated cells (scATACseq, 24 h). Motifs with high chromatin access in the clusters with high nucleosome signal (see panel S3D) corresponding to chromatin state 5 (DMSO: left) and chromatin state 6 (AZD1775: right) are shown. Related to Fig. 3A-B.

**C.** TF motif activity score visualized for TFs shown in B on 10 h DMSO scRNA-seq UMAP, as in Fig. 3A.

**D**. Nucleosome signal visualized as violin plot from scATAC 24 h and 10 h data filtered based on other quality metrics (See methods). Grey: DMSO treated cells. Green: AZD1775-treated cells. Final analysis was limited to cells with signal <10.

**E.** Quality metrics compared between chromatin state clusters identified from scATAC 24 h data. Initial clustering based on the neighborhood graph obtained from LSI-reduced scATAC- seq signal resulted in 6 (DMSO) and 7 (AZD1775) clusters. From both treatments, cells with lowest peak region fragments were removed from further downstream analysis. Condensed chromatin state corresponds to clusters 5 (DMSO) and 6 (AZD1775).

**F.** scRNA-seq transcriptome based UMAP visualizations (top) of Nalm-6 cells treated with DMSO or AZD1775 are shown. Colors correspond to matching RS4;11 cell state. NA: cells with prediction score < 0.5. Cell states 6 and 7 correspond to highest p53 activity. Comparison of cell state proportions based on scRNA-seq analysis of RS4;11 and Nalm-6 cell lines is shown (bottom).

**G.** Immunoblot analysis of γH2AX and pS10-H3 in response to AZD1775 and after drug washout (WO) for 12 and 24 hrs in RS4;11 and Nalm-6 cells, respectively. Histone 3 serves as a loading control.

#### Figure S4

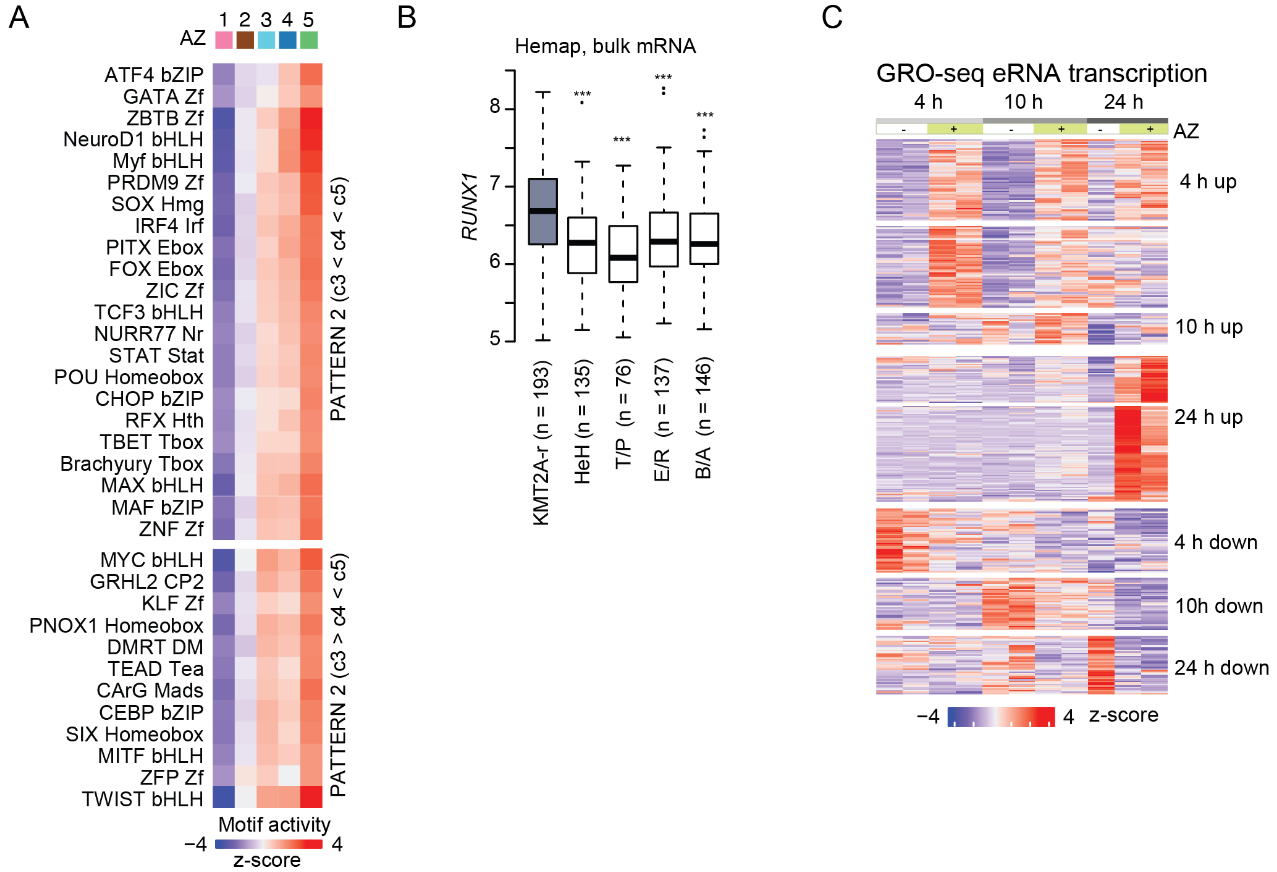


**Fig. S4 related to Fig. 4.**

**A.** Heatmap showing TF motif activity (scATAC, 24 h) across chromatin states in AZD1775 treated cells for additional TF motifs in Pattern 2 (see Fig. 4A).

**B.** Boxplots comparing *RUNX1* gene expression in B-ALL subtypes based on HEMAP data from diagnostic patient samples. Wilcoxon test p-value from KMT2A-r vs each other subtype is indicated (*** : p-value < 0.001).

**C.** Heatmap illustrating the magnitude and direction of changes in the GRO-seq eRNA signal (z-score; tones of red indicate high level, tones of blue low level). Top enriched TF motifs for the corresponding DNA sequences are listed in Table S4.

**Figure S5**

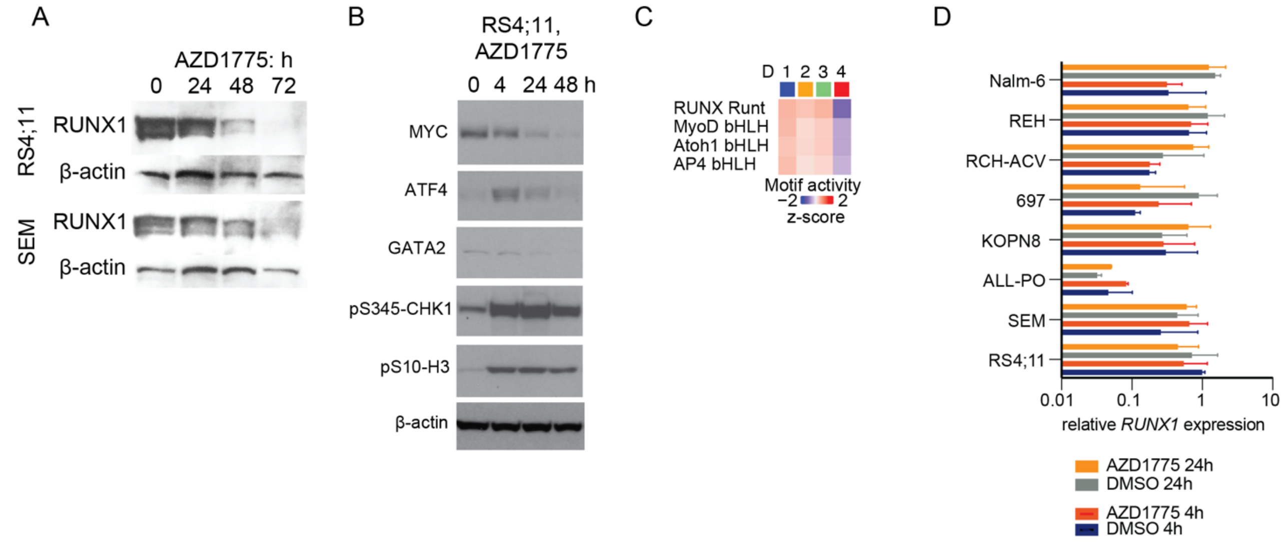


**Fig. S5 related to Fig. 5:**

**A.** RUNX1 protein levels in RS4;11 and SEM treated with AZD1775 or DMSO for indicated time points were analysed by immunoblotting. β-actin served as a loading control.

**B.** Whole cell lysates of RS4;11 cells treated with AZD1775 for the indicated time points and immunoblotted with antibodies against, MYC, ATF4, GATA2, pCHK1, pS10-H3. b-actin is a loading control.

**C.** Motifs with similar profile to RUNX1 (refer to Fig. 4A) visualized as heatmap from DMSO (D) treated cells (24 h). Chromatin states identified from DMSO scATAC-seq data are shown (see Fig. S3A).

**D.** *RUNX1* mRNA expression levels at 4 h and 24 h treatment (DMSO compared to AZD1775) was analysed by quantitative PCR (qPCR). Relative *RUNX1* expression levels were adjusted to the *GAPDH* expression.

#### Figure S6

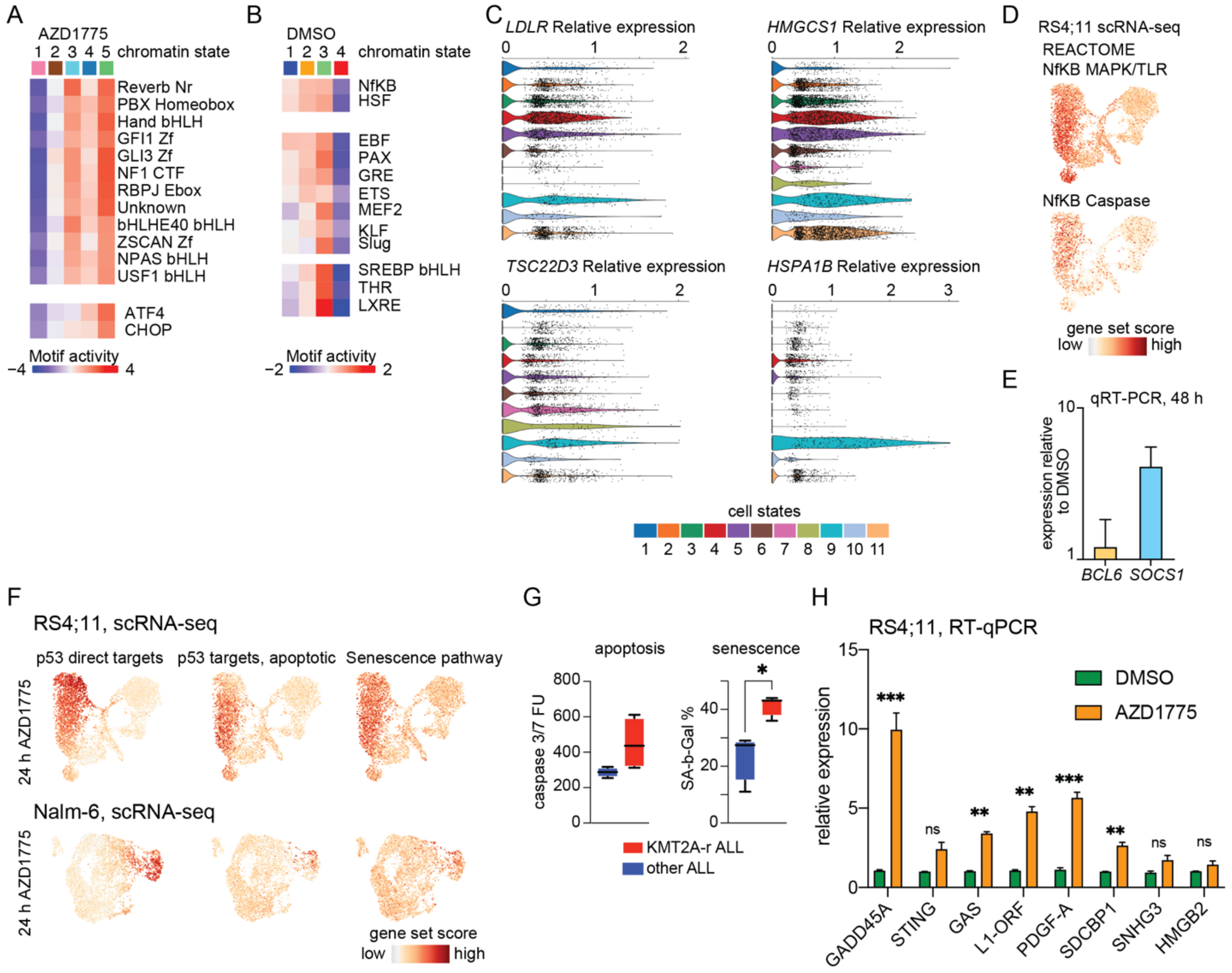


**Fig. S6 related to Fig. 6.**

**A-B**. TF motif access across 24 h AZD1775 (A) and DMSO (B) chromatin states is shown in the heatmap for TFs with high activity in chromatin state 5 (see Fig. 6A).

**C.** *LDLR, HMGCS1, TSC22D3* and *HSPA1B* expression level is shown from scRNAseq 24 h as violin plots comparing cell states 1-11. The track color corresponds to cell state annotation.

**D.** Gene set score for public gene sets related to NfKB MAPK/TLR and NfKB caspase targets visualized on the UMAPs in 24 h AZD1775 treated cell sample. Red color tones indicate higher gene set score.

**E.** Quantitative reverse transcription PCR (qRT-PCR) analysis of *BCL6* and *SOCS1* mRNA levels at 48 h AZD1775 treatment adjusted to the *GAPDH* mRNA, relative to the DMSO control is shown.

**F**. Gene set score for P53 direct target genes is visualized on the scRNA-seq UMAP for RS4;11 (top) and Nalm-6 cells (bottom) treated with AZD1175 for 24 h.

**G.** Box-plots comparing Caspase3/7 activity in response to AZD1775 treatment in KMT2A-r and non-KMT2A-r ALL cell lines. Box plot visualization of the percentage of positive cells. Senescence associated-β-Galactosidase (SA-β-Gal)-positive cells following 72 hr of AZD1775 treatment and 10 days drug wash-out comparing KMT2A-r and non-KMT2A-r ALL cell lines. * indicates p=0.0167 determined using t-test. Data are represented as mean ± SD.

**H**. qPCR analysis of senescence-associated secretory pathway (SASP) genes. mRNA expression levels of the indicated genes at 48 h of AZD1775 treatment is shown adjusted to the *GAPDH* mRNA and relative to the DMSO control. ** denotes $p<0.01$ and *** denotes $p<0.001$ determined using Student's t-test. Data are represented as mean ± SD.

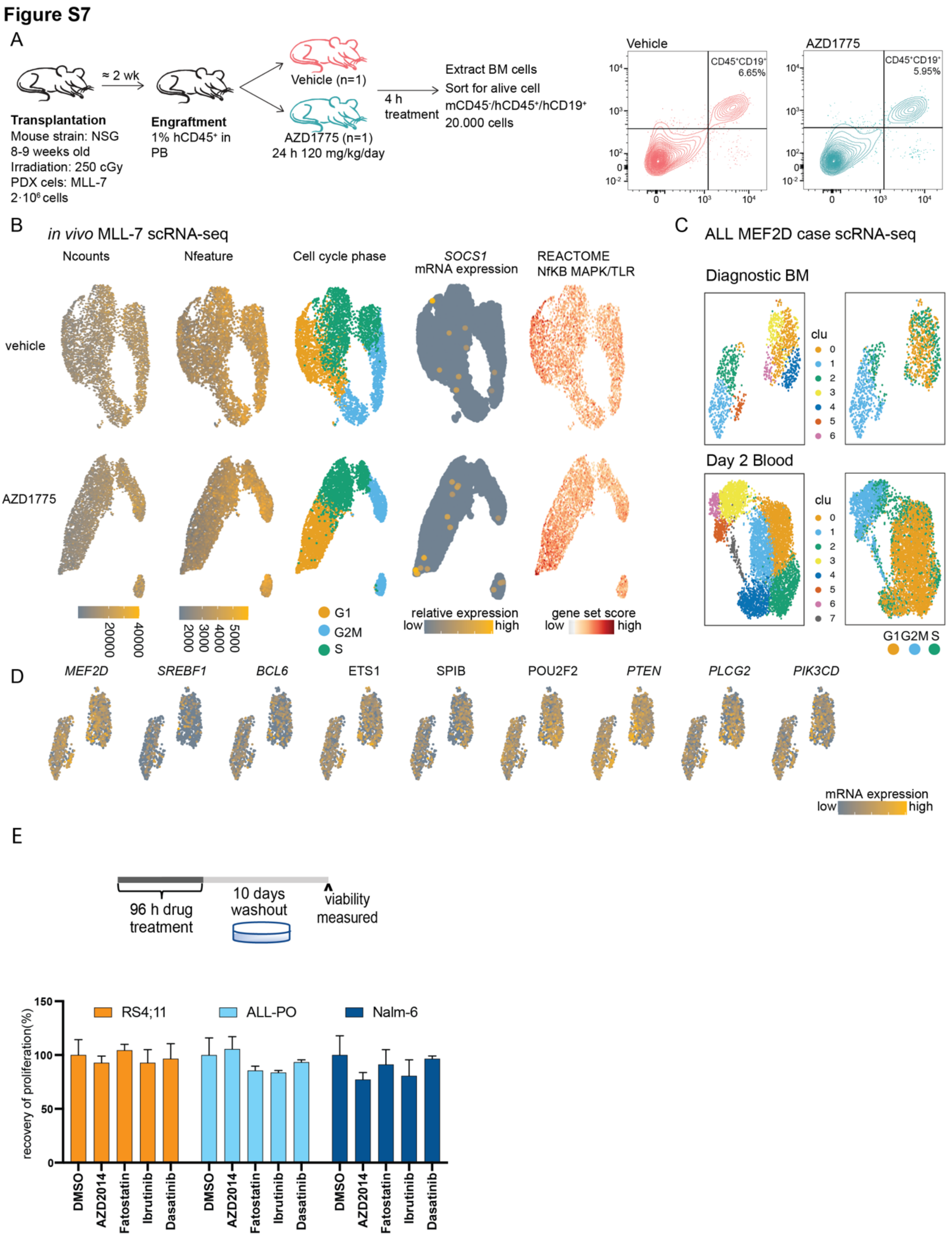


**Fig. S7 related to Fig. 7.**

**A**. Schematic representation of the experimental plan depicting the treatment arms and downstream analysis. After 28 hours and two doses of 120 mg/kg AZD1775 animals were sacrificed and the cells were analysed by flow cytometry and hCD45+hCD19+ cells were sorted for scRNA-seq.

**B.** scRNA-seq UMAPs from DMSO (top) and AZD1775-treated (bottom) primary MLL-7 cells. Quality metrics (left; N counts: number of counts, N feature: number of genes), assigned cell cycle status (middle) and mRNA level for *SOCS1* gene visualized (brighter color tones correspond to higher level). Right: Gene set score for public gene sets related to NfKB MAPK/TLR (Red color tones indicate higher values.)

**C-D.** Primary MEF2D-fusion ALL case treated with standard chemotherapy (top: diagnostic bone marrow below: day 2 blood). Clustering, cell cycle state and mRNA levels for BCR-related genes are visualized as in B.

**E** Upper panel: overview of the experimental setup. Cells were treated for two days with indicated drugs and allowed to recover for an additional 10 days without the drugs. Lower panel: recovery of cell proliferation following drug removal. Regrowth was assessed using Alamar Blue stainings. Data are represented as mean ± SD.
